# Imbalance of Neuregulin1-ErbB2/3 signaling underlies altered myelin homeostasis in models of Charcot-Marie-Tooth disease type 4H

**DOI:** 10.1101/2022.01.20.477077

**Authors:** Lara El-Bazzal, Adeline Ghata, Clothilde Estève, Patrice Quintana, Nathalie Roeckel-Trévisiol, Frédérique Lembo, Nicolas Lenfant, Andre Mégarbané, Jean-Paul Borg, Nicolas Lévy, Marc Bartoli, Yannick Poitelon, Pierre L. Roubertoux, Valérie Delague, Nathalie Bernard-Marissal

## Abstract

Charcot Marie Tooth disease (CMT) is one of the most common inherited neurological disorders, affecting either axons from the motor and/or sensory neurons or Schwann cells (SC) of the peripheral nervous system, and caused by more than 100 genes. We previously identified mutations in *FGD4*, as responsible for CMT4H, an autosomal recessive demyelinating form of CMT. *FGD4* encodes FRABIN a GDP/GTP nucleotide exchange factor (GEF), particularly for the small GTPase cdc42. Remarkably, nerves from patients with CMT4H display excessive redundant myelin called outfoldings that arise from focal hypermyelination, suggesting that FRABIN could play a role in the control of PNS myelination. To gain insights into the role of *FGD4*/FRABIN in SC myelination, we generated a knock-out mouse model, with conditional ablation of *fgd4* in SC. We showed that the specific deletion of FRABIN in SCs leads to aberrant myelination *in vitro*, in dorsal root ganglion (DRG)/SCs cocultures as well *in vivo*, in distal sciatic nerves. We observed that those myelination defects are related to an upregulation of some interactors of the NRG1 type III/ErbB2/3 signaling pathway, which is known to ensure a proper level of myelination in the PNS. Based on a yeast two-hybrid screen, we identified SNX3 as a new partner of FRABIN, which is involved in the regulation of endocytic trafficking. Interestingly, we showed that loss of FRABIN impairs endocytic trafficking which may contribute to the defective NRG1 type III/ErbB2/3 signaling and myelination. Finally, we showed that the reestablishment of proper levels of the NRG1 type III/ErbB2/3 pathway using Niacin treatment reduces myelin outfoldings in nerves of CMT4H mice. Overall, our work reveals a new role of FRABIN in the regulation of NRG1 type III/ErbB2/3 NRG1signaling and myelination and opens future therapeutic strategies based on the modulation of NRG1 type III/ErbB2/3 NRG1to reduce CMT4H pathology and more generally others demyelinating CMT.

## Introduction

Charcot-Marie-Tooth (CMT) disease also known as Hereditary Motor and Sensory Neuropathy (HMSN) is one of the most common inherited groups of neurological diseases, with an overall prevalence of about 1/2500 (Skre 1974). Clinically, CMT diseases are characterized by progressive muscular weakness starting at the distal extremities, pes cavus deformity, loss of deep tendon reflexes, associated with mild to moderate distal sensory loss (Harding and Thomas 1980). Two main subgroups can be defined based on electrophysiological and histopathological characteristics: the demyelinating form (CMT1) resulting from primary damage of myelinating Schwann cells and the axonal form (CMT2) originating on the neuronal/axonal side. Patients affected with CMT1 have reduced nerve-conduction velocities (NCV, ≤38m/s), whereas patients affected with CMT2 show slightly reduced to normal NCVs but reduced amplitudes (≥38m/s) (Pareyson, Scaioli et al. 2006). In addition, CMT disease is characterized by extensive clinical and genetic heterogeneity, with around 100 genes identified to date and mutations segregating following all modes of inheritance (Pipis, Rossor et al. 2019). Autosomal recessive demyelinating forms of CMT (CMT4) are less frequent, usually of earlier onset and more severe, than the autosomal dominant CMT subtypes (CMT1), leading to a higher frequency of wheelchair-dependency in the life course (Berciano and Combarros 2003, Baets, Deconinck et al. 2011).

We identified in 2007, FGD4/FRABIN as a causative gene of the CMT4H, a rare autosomal recessive demyelinating form of CMT (Delague, Jacquier et al. 2007). The disease is characterized by typical findings of CMT, such as distal amyotrophy and foot deformities, with early-onset and slow progression. Motor and Sensory Nerve Conduction Velocities (NCVs) are strongly reduced in all patients, but there is a strong inter-and intra-familial variability in the severity of the disease. The diagnosis is established by the presence of biallelic pathogenic variants in *FGD4*. More than 30 mutations are described to date in *FGD4*, most of them resulting in a loss of the protein or a non-functional truncated protein (Delague, Jacquier et al. 2007, Baudot, Esteve et al. 2012, Boubaker, Hsairi-Guidara et al. 2013). *FGD4* encodes FRABIN, a Guanine Exchange Factor (GEF), carrying a Dbl Homology domain responsible for GDP/GTP exchange on the small RhoGTPases and more particularly Cdc42 and Rac1 (Umikawa, Obaishi et al. 1999, Ono, Nakanishi et al. 2000). Remarkably, nerves from patients affected with CMT4H display characteristic myelin abnormalities defined as myelin outfoldings (i.e., redundant loops of myelin), which arise from aberrant focal hypermyelination. In the Peripheral Nervous System (PNS), the myelination process is mainly regulated by the NRG1 type III/ErbB2/3 pathway. Notably, the amount of axonal NRG1 type III, and its downstream signaling via the activation of the ErbB2/3 receptors, determines the thickness of the myelin sheath (Michailov, Sereda et al. 2004, Taveggia, Zanazzi et al. 2005). In consequence, this pathway has to be tightly regulated to avoid improper myelination. Previous work has reported that myelin outfoldings might arise from an enhanced NRG1 type III/ErbB2/3 signaling, as well as the overactivation of the downstream effectors AKT/mTOR (Goebbels, Oltrogge et al. 2012, Bolino, Piguet et al. 2016, Domenech-Estevez, Baloui et al. 2016). A growing body of work suggests that the dysregulation of ErbB receptors’ trafficking may be the cause of NRG1 type III/ErbB2/3 signaling’s impairment in Schwann cells (SCs) and a common pathogenic mechanism for several CMT4 subtypes, such as CMT4B1, CMT4B2, and CMT4C (Lee, Chin et al. 2017, Markworth, Bahr et al. 2021).

The implication of FRABIN in the regulation of PNS myelination and the above-mentioned signaling pathways remains poorly studied. Interestingly, in addition to its Dbl domain, responsible for the GEF activity toward Cdc42 and Rac1, FRABIN contains several other functional domains, including an N-terminal F-actin binding domain as well as two Pleckstrin Homology (PH) and one FYVE domains, which are implicated in binding with Phosphoinositides (PIs). PIs are lipid regulators of endocytic trafficking (Cullen and Carlton 2012). Previous work by Horn et al (Horn, Baumann et al. 2012) showed an impairment of endocytosis in the absence of FRABIN, but its link to aberrant hypermyelination has not been studied so far.

We hereby provide evidence that FRABIN is required for proper myelination of the PNS by regulating NRG1 type III/ErbB2/3 signaling in both *in vitro* and *in vivo* models of CMT4H. In particular, we show that the specific deletion of FRABIN in SCs leads to aberrant myelination *in vitro*, in dorsal root ganglion (DRG)/SCs cocultures, and *in vivo*, in distal sciatic nerves. We demonstrate that the myelination defects in CMT4H are related to impaired NRG-1 type III/ErbB2/3 signaling and defective endocytic trafficking. Finally, we show that the reestablishment of proper levels of the different actors of the NRG1 type III/ErbB2/3 pathway using Niacin treatment reduces myelin outfoldings in nerves of CMT4H mice.

## Material and Methods

### Animals

To generate a conditional *Fgd4*-null allele, we first generated a mouse with a floxed allele by flanking *Fgd4* exon 4 with *lox-P* sites. Its excision, by crossing with a transgenic mouse expressing the Cre-recombinase, introduces a frameshift and is predicted to generate a premature stop codon in exon 5. Mice heterozygous for the *Fgd4* floxed allele (*Fgd4*^*fl/+*^) have been developed on a pure C57BL/6N background in collaboration with “La Clinique de la Souris” (ICS; Strasbourg; http://www.ics-mci.fr) by performing homologous recombination in ES-derived from C57BL/6N mouse. The *Fgd4*^*fl/+*^ line was maintained on a pure C57BL/6N background (Taconic Biosciences A/S, Danemark). Genotyping of *Fgd4*^*fl/+*^ mice was performed by PCR on DNA isolated from tail biopsies using Direct PCR lysis reagent (Viagen Biotech Inc., USA), following the manufacturer’s instructions, using the primers flox-LF (5’-CGAACCCTTAGCGATCTGTT-3’) and flox-LR (5’-TTTTCCTAGCTGGCGTGTTT-3’).

To obtain an allele with specific deletion of *Fgd4* in SCs, we crossed *Fgd4*^*fl/+*^ mice with transgenic mice expressing the Cre-recombinase under a promoter-specific of SCs (B6N.FVB-Tg(Mpz-cre)26Mes/J), “so-called P0-Cre mice”, available at the Jackson Laboratories (#017927) (Feltri, D’Antonio et al. 1999). We obtained *Fgd4*^*fl/+*^; *P0-Cre*,, so-called *Fgd4*^*SC-/+*^ that we backcrossed for at least 10 generations on C57BL/6N (Taconic) background. The P0-Cre transgene was detected by PCR using the following primers: P0Cre-F (5’-CCACCACCTCTCCATTGCAC-3’) and P0Cre-R (5’-GCTGGCCCAAATGTTGCTGG-3’). For ubiquitous deletion of *Fgd4*, (*Fgd4*^*fl/+*^; *CMV-Cre*, mice, so-called *Fgd4*^*-/+*^), were generated by crossing the floxed *Fgd4*^*fl/+*^ line with transgenic mice expressing the Cre-recombinase under the CMV promoter (B6.C-Tg (CMV-cre)1Cgn/J), available at the Jackson Laboratories (#006054). The CMV-Cre transgene was detected by PCR using the following primers: CMVCre-F (5’-AGGTTCGTTCACTCATGGA-3’) and CMVCre-R (5’**-**TCGACCAGTTTAGTTACCC-3’). The excision of the exon 4 in the *Fgd4*^*-/+*^ mice was detected using the primers flox-LF (5’-CGAACCCTTAGCGATCTGTT-3’) and flox-ER (5’-CAAGCCTCAGCTTCACTTCC-3’). Animals were housed in an animal facility with a 12 h light / 12 h dark environment, and *ad libitum* access to water and a normal diet. All experiments were done in accordance with a national appointed ethical committee for animal experimentation (Ministère de l’Education Nationale, de l’Enseignement Supérieur et de la Recherche; Authorization N°2019062110352453 v4).

### RT-PCR

Total RNA was isolated from Wild-Type (WT) and *Fgd4*^*SC-/-*^ mouse sciatic nerves using the Purelink RNA Minikit (#12183018A, Thermofisher Scientific, USA), following the manufacturer’s protocol. cDNA was generated using the SuperScript III one-step RT-PCR (#12574018, ThermoFisher Scientific, USA) and random hexamers (#48190011, Thermofisher Scientific, USA). The deletion of exon 4 was detected by the amplification, from cDNA, of a fragment between exons 3 and 7 of the Fgd4 transcript, using the following primers (Fgd4-mouse-3F: 5’-GAGTCTAATCCGGCCCCTAC-3’and Fgd4-mouse-7R: 5’-AAGGAATGGCGCCAACTTTT-3’).

### Behavioural gait test

For the gait experiment, sex and age-matched F*gd4*^*SC-/-*^ and WT littermates’ mice were monitored at 6 (n=9 WT and n=9 F*gd4*^*SC-/-*^), 12 (n=12 WT and n=13 F*gd4*^*SC-/-*^) and 18 (n=11 WT and n=13 F*gd4*^*SC-/-*^) months-old. Gait was analyzed during spontaneous walk using an automated gait analysis system (Gaitlab, Viewpoint, France). Before recording footprints, mice were acclimated and trained to walk on the Gait system for two days. During the testing session, a minimum of 2-3 completed runs were collected. We focused the analysis on intensity-based parameters, paw-size as well as gait/posture, as previously described in (Bernard-Marissal, van Hameren et al. 2019). Animals that did not complete at least 2 successful runs (stalling or reversing during gait,…) were removed from the study.

### Niacin treatment

One-month-old *Fgd4*^*SC-/-*^ mice were daily injected with saline solution or Niacin (60mg/Kg) for 8 weeks (n=4 saline-treated animals and 5 Niacin-treated animals). Nicotinic acid (#PHR1276, Niacin, Sigma-Aldrich, USA) was diluted in sterile saline solution and was daily injected intraperitoneally at a dose of 60 mg/kg.

### Electron microscopy and morphometric analysis

The sciatic nerves were dissected and fixed in 2% ParaFormaldehyde (PFA) and 2.5% Glutaraldehyde in 0.1M cacodylate buffer for 2 hours. The next day, the nerves were washed three times in 0.1M cacodylate buffer and post-fixed in buffered 1% OsO4 for one hour. After washes in distilled water, the samples were contrasted in aqueous 1% uranyl acetate. Samples were then dehydrated in graded series of ethanol baths (30 minutes each) and infiltrated with epon resin in ethanol (1:3, 2:2, 3:1) for 2 hours for each, and finally in pure resin overnight. The next day the nerves were embedded in fresh pure epon resin and cured for 48h at 60°C. 500 nm semi-thin and 70 nm ultra-thin sections were performed on a Leica UCT Ultramicrotome (Leica, Austria). Semi-thin sections were stained with toluidine blue and ultrathin sections were deposited on formvar-coated slot grids. The grids were contrasted using uranyl acetate (10 minutes) and lead citrate (5 minutes) and observed using an FEI Tecnai G2 at 200 KeV. The acquisition was performed on a Veleta camera (Olympus, Japan). The proportion of fibers having out- and infoldings was counted on ten fields corresponding to a range of 600-800 fibers for each sciatic nerve of at least three animals.

To perform g-ratio analysis, digitalized images of fiber semithin sections of the sciatic nerves were obtained with a 100× objective of a phase-contrast microscope (BX59, Olympus). At least ten images from three different animals per genotype at 3, 6, and 18 month-old were acquired. G-ratio analysis was performed on micrographs using Image J (National Institutes of Health, Bethesda, MD) plug-in (g-ratio calculator) developed in collaboration with the cellular imaging facility of the University of Lausanne and available at http://cifweb.unil.ch, as previously described in Arnaud E 2009 (Arnaud, Zenker et al. 2009).

### Immunohistochemistry

Mice were sacrificed with an overdose of CO_2_. Gastrocnemius muscles were dissected out and fixed in 4 % PFA for 30 min. Samples were then incubated in 25 % sucrose solution at 4 °C for 24 h. Tissues were embedded in the Optimal Cutting Temperature compound (#3801480, Leica, USA) and stored at -80 °C before processing. Embedded muscles were then cut to twenty-five μm-thick sections, mounted on slides coated with Superfrost Plus (#LSFPLUS, THERMO SCIENTIFIC MENZEL, USA). Muscle sections were incubated overnight at 4 °C in blocking solution (2 % BSA, 10 % Normal Goat Serum, 0.1 % Triton and PBS) with chicken anti-NFM (#822701, BioLegend, USA, previously #PCK-593P, Covance, USA). Sections were next incubated with Goat anti-Chicken IgY (H+L) Secondary Antibody, Alexa Fluor 488, (#ab150169, Abcam,UK,)USA) and *α*-Bungarotoxin, Alexa Fluor™ 555 conjugate (#B35451, thermofisher Scientific, USA), (1:500), and mounted in Duolink mounting medium (#DU082040, Sigma-Aldrich, USA). Neuromuscular junctions were imaged on a Zeiss ApoTome.2 microscope (Zeiss, Germany) equipped with an AxioCam MRm camera.

### Primary cell culture

#### Schwann cells culture

Primary rat SCs were prepared from the sciatic nerve of postnatal day 3 pups. SCs were maintained in DMEM (#41965, Thermofischer Scientific, USA), 2 mM L-glutamine (#25030024, Thermofisher Scientific, USA), 10 % FBS (#10270098, Thermofisher Scientific, USA), 2 μM forskolin (#344270, Merck, Germany) and 2 ng/ml recombinant human NRG1-β 1 (#396-HB, R&D Systems, USA) as described in Poitelon *et al*. (Poitelon and Feltri 2018). SCs were not used beyond the fourth passage.

#### Dorsal root ganglia (DRG)/SCs co-cultures

Mouse DRG neurons were isolated after dissecting the spinal cord of embryonic day 13.5 (E13.5) embryos and seeded on 12 mm glass coverslips, as described (Poitelon and Feltri 2018). In details, after isolation, DRG were dissociated and then incubated in 0.25 % Trypsin solution (#25200056, Thermofisher Scientific, USA) for 45 min at 37 °C. DRGs were mechanically dissociated and ∼40000 cells were seeded in matrigel (#356234, Corning, USA), USA) coated coverslips in C-medium, composed ofMEM medium (#11090081, Thermofisher Scientific, USA), 2 mM L-glutamine (#25030024, Thermofisher Scientific, USA), 10 % FBS (#10270098, Thermofisher Scientific), 4 mg/ml D-glucose (#G5146 (Sigma-Aldrich, USA), 50 ng/ml NGF (#N6009, Sigma-Aldrich, USA). After 24 h, the C-medium was replaced by NB medium:Neurobasal medium (#21103049, Thermofisher Scientific, USA) 4 g/l D-glucose (#G5146, (Sigma-Aldrich, USA), 2 mM L-glutamine (#25030024, Thermofisher Scientific, USA), 50 ng/ml NGF (#N6009, Sigma-Aldrich), and B27 supplement 1 × (50X, #17504044, Thermofisher Scientific, USA) and changed every two days. After 5 days of NB medium, myelination was induced by adding 50 µg/ml ascorbic acid (#A0278, Sigma-Aldrich, USA) to the c-medium, for 12 days.

### Drugs

Co-cultures were treated using Nicotinic acid (Niacin,#PHR1276, Sigma-Aldrich, USA) diluted in MEM (#11090081, Thermofisher Scientific, USA), at a final concentration of 5 mM. Endosidin-2 (#SML1681-5mg, Sigma-Aldrich, USA) was diluted in MEM at a final concentration of 1 µM. Niacin or Endosidin-2 were added concomitantly to ascorbic acid at day 6 of the co-culture.

### Lentivirus infection

Fgd4/FRABIN was either overexpressed or downregulated in WT DRG/SC cocultures or in rat primary SC using Lentivirus vectors (Lv) designed and produced by the Vector builder company (https://en.vectorbuilder.com, USA). For overexpression, we designed control Lv (Lv-control) expressing EGFP under the CMV promoter and Lv allowing the expression of mouse Fgd4 (Lv-Fgd4) under the CMV promoter. For the knockdown experiments, we used an Lv expressing an shRNA against rat *Fgd4* previously validated in Horn et al. 2012 (Horn, Baumann et al. 2012)(targeted sequence: 5’-GAAGAAGAGGATATTGTA-3’) referred to as LvSHFgd4-GFP. Lv-Shcontrol-GFP expressing an shRNA scramble, randomly produced, was provided by Vectorbuilder (#VB151023-10034, Vectorbuilder, USA). Both Lv vectors express EGFP under the CMV promoter, in addition to the shRNA, allowing tracing of the infected cells.

We used the same strategy to knockdown the *Snx3* gene in DRG/SC cocultures or in primary SCs. Here, we used Lv expressing two shRNAs targeting *Snx3* (Sh-0: 5’-GCCCAGAATGAACGTTGTCTT-3’; Sh-3: 5’-AGAGAGAGCAAGGTTGTAGTT-3’), both provided by Sigma-Aldrich (USA). Cells (cocultures and primary SCs) were infected with the described Lv at a dose of 15 TU/cell 24 h after plating.

### Transferrin assay

Rat primary SCs were plated at a density of 50 000 cells per well and infected 24h later with Lv-SHcontrol-GFP or SHFgd4-GFP (15 TU/cell). Two days after the infection, cells were incubated with pHrodo-red Transferrin conjugate (#P35376, Thermofischer Scientific, USA) at a dilution of 50µg/ml, for 30 min on ice, followed by 15, 30 or 45 min of incubation at 37 °C. After one PBS wash, cells were fixed in 4% PFA for 15 min. Samples were then mounted in Duolink mounting medium (#DU082040, Sigma-Aldrich, USA) and pHrodo-TF fluorescence intensities were visualized and captured on a Zeiss ApoTome.2 microscope (Zeiss, Germany) equipped with an AxioCam MRm camera, with a 20 X objective. The ntegrated density of fluorescence signal was analyzed only in the infected cells (*i*.*e*., GFP+) using ImageJ software.

### Cell lines and transfection

HEK293 (Human embryonic kidney, # CRL-1573) and S16 rat SC (#CRL-2941) lines were both provided by ATCC (Manassas, Virginie, USA). HEK293 and S16 cells were cultured in DMEM (#41965, Thermofisher Scientific, USA) complemented with 10 % of FBS (#15000-036, Thermofisher, USA) and 1 % of penicillin/streptomycin (#15070063, Thermofisher Scientific, USA). To overexpress hFGD4/FRABIN in HEK293 cells, the pEx-FGD4-His-V5 vector was generated by Gateway cloning technology (Thermofisher, USA). Transfection experiments were performed using promofectin reagent (#PK-CT-2000-50, PromoCell GmbH, Germany). Briefly, 300000 cells were seeded and transfected with 4µg of plasmid following the manufacturer’s recommendation. For immunoprecipitation, cells were harvested 72 h post-transfection.

### Plasmids

The plasmids used for the yeast-two hybrid experiment (pENTR-FL-hFGD4) and transfection studies (pEx-FGD4-His-V5) were generated using Invitrogen Gateway cloning technology (ThermoFischer Scientific, USA) following the manufacturer’s instructions. Briefly, the entry plasmid (i.e., pENTR-FL-hFGD4) was generated after the BP recombination reaction between the *att*B-flanked hFGD4 fragment and the *att*P-containing donor vector pDONR221 (#12536017, ThermoScientific, USA) using the BP clonase enzyme kit (#11789013, ThermoScientific, USA). The *att*B-flanked hFGD4 fragment was produced by PCR using the “Expand High Fidelity Plus PCR System” (#3300226001, Roche, Switzerland) and the following primers: FGD4-GATEWAY-1F:5’GGGGACAAGTTTGTACAAAAAAGCAGGCTTCGAGGAAATTAAACCTGCCTCTGC3’ and FGD4-GATEWAY-1R: 5’GGGGACCACTTTGTACAAGAAAGCTGGGTTCAGCATTCTGATTTTTTCTTAGG3’. The expression vector pEx-FGD4-His-V5 was generated following an LR recombination between the attL containing entry vector (pENTR-FL-hFGD4) and the attR destination vector (pDEST40, #12274015, ThermoScientific, USA) using the LR Clonase enzyme mix (#11791019, ThermoScientific, USA).

### RNA sequencing

RNA-Sequencing and bioinformatics analysis was performed by the Genomics and Bioinformatics facility (GBiM) from the U 1251/Marseille Medical Genetics lab.

mRNA sequencing (mRNA-Seq) was performed, in duplicate, on total RNA samples extracted from DRG/SCs cocultures derived from either conditional knock-out (*Fgd4*^*SC-/-*^) or Wild-Type littermates’ embryos (4 samples in total). Total RNA was extracted from each condition (2 coverslips of control Fgd4^fl/fl^ and *Fgd4*^*SC-/-*^ DRG/SC cocultures, in two replicates), using Purelink silica-membrane, anion exchange resin, spin-column kits, following the manufacturer’s recommendations (Purelink RNA Minikit, #12183018A, ThermoFisher Scientific, USA). Before sequencing, the quality of total RNA samples was assessed by using the Agilent Bioanalyzer (Agilent Technologies, USA): only RNAs with RNA Integrity Numbers (RIN) above 8 were deemed suitable for sequencing and used for library preparation. For each sample, a library of stranded mRNA was prepared from 500 ng of total RNA after capture of RNA species with poly-A tail poly(A), using the Kapa mRNA HyperPrep kit (Roche, Switzerland), following the manufacturer’s instructions. The quality and profile of individual libraries have been quantified and visualized using Qubit™ and the Agilent Bioanalyzer dsDNA High Sensibility Kit (Agilent Technologies, USA) respectively. Indexed libraries were pooled and sequenced (2*75 bp paired-end sequencing) on an Illumina NextSeq 500 platform (Illumina, USA).

### Data processing and differential gene expression analysis (DGE)

The quality of sequencing reads was assessed using fastQC v0.11.5 (https://www.bioinformatics.babraham.ac.uk/projects/fastqc/). Raw sequencing reads were mapped to the mouse reference genome (Mus Musculus genome assembly GRCm38 (mm10) using STAR v2.5.3a (Dobin and Gingeras 2015), and bam files were indexed and sorted using Sambamba v0.6.6 (Tarasov, Vilella et al. 2015). After mapping, the number of reads per feature (GENCODE v34 annotations) was determined using Stringtie v1.3.1c (Pertea, Pertea et al. 2015). Differential gene expression analysis was performed using a Wald test thanks to the DESeq2 package (Love, Huber et al. 2014). P-values were adjusted for multiple testing using the method by Benjamini and Hochberg (Benjamini and Hochberg 1995). Only transcripts with an adjusted p-value (FDR, False Discovery Rate) below 0·05 were considered as significantly differentially expressed. Relative expressions of the most variable features between samples have been plotted as heatmaps using the Pheatmap R package. Differentially Expressed Genes (DEGs) were visualized in the form of volcano plots, representing the Log2FC (log2 of the expression fold change) and the adjusted p-value, prepared using the EnhancedVolcano R package. Enrichment (Gene set enrichment Analysis, GSEA) and overrepresentation (Singular Enrichment Analysis, SEA) of GO term annotation in the DEGs were performed using a Mann-Whitney test and hypergeometric test respectively, using enrichGO from the R-package clusterProfiler (v3.10.15). For functional Gene Ontology (GO) annotation, we selected the differentially expressed genes, with a statistical significance of padj value < 0.05 (SEA). Pathway analysis was conducted, using Reactome, which use a hypergeometric test with an adjustment of the p-value (BH) to detect pathway, enriched in the dataset of genes (DEG) from the top10 GO BP terms.

### Immunocytochemistry

Co-cultures were fixed with 4% PFA, washed in PBS, and then permeabilized for 5 minutes with methanol. After two PBS washes, cells were incubated 1-2 h in a blocking solution (20% fetal bovine serum, 1% bovine serum albumin, 0.01% Triton in 1 X PBS) at room temperature. Cells were then incubated with the following primary antibodies diluted in the incubation solution overnight: chicken anti-neurofilament NF-M (1:1000) (#822701, BioLegend, USA, previously #PCK-593P, Covance), rat anti-MBP (1:300) (#MAB386, Merck-Millipore, Germany). After two washes in PBS, cells were incubated with one of the following secondary antibodies: Donkey Anti-Rat IgG H&L (#150154, Alexa Fluor 555, abcam, UK) (1:1000) and (#A21449, Goat Anti-Chicken IgY H&L (Alexa Fluor 647, Invitrogen, USA) (1: 1000). Coverslips were then rinsed twice in PBS and mounted in a duolink mounting medium (#DU082040, Sigma-Aldrich, USA) for microscope analysis. Fluorescence images were captured with a Zeiss ApoTome.2 microscope (Zeiss, Germany) equipped with an AxioCam MRm camera. Images were captured and merged with the ZEN software (Zeiss, Germany) and were treated using ImageJ software.

### Analysis of myelination

Around 10 fields per coverslip were randomly acquired with a Zeiss ApoTome.2 microscope (Zeiss, Germany) at a 20X objective. Myelin abnormalities were defined as an excessive redundant or abnormal thickening of myelin visualized along with MBP positive segments and quantified in ∼ 150-160 myelinated segments for each coculture and at least 3 cocultures. Results were expressed as a percentage of MBP positive segments showing myelin abnormalities on the total of MBP positive fibers.

### Protein extraction and Immunoblotting

Co-cultures or nerve samples were lysed in RIPA buffer supplemented with a protease/phosphatase inhibitor cocktail (#78442, ThermoFisher Scientific, USA). The lysate was passed through an 18–21-gauge needle and submitted to sonication using the Bioruptor VCD-200 (Diagenode, Belgium). After centrifugation for 10 min at 20,000 x g at 4 °C, the supernatant was removed and protein concentration was measured by use of a Bicinchoninic acid (BCA) solution (#B9643-1LSigma-aldrich, USA) coupled to copper II sulfate solution (#C2284-25ml, Sigma-aldrich, USA) following the manufacturer’s recommandations. 40 μg of samples’ proteins were loaded on a Precast NuPage 4-12 % Bis-Tris gels (Thermofisher, USA) and transferred onto a nitrocellulose membrane (GE healthcare life science, Germany). The membrane was then blocked by incubation in blocking buffer (Intercept-blocking buffer, Licor). The membrane was then incubated overnight with the following primary antibodies: mouse anti-Neuregulin1α/β1/2 (D10) (1:500) (#sc-393009; Santa Cruz Biotechnology, USA), rabbit anti-phospho-Akt (Ser473) (D9E) (1:2000) (#4060, Cell Signaling Technology, USA), rabbit anti-Akt(pan) (C67E7) (1:1000) (#4691, Cell Signaling Technology, USA), rabbit anti-HER2/ErbB2 (29D8) (1:1000) (#2165, Cell Signaling Technology, USA), rabbit anti-phospho-HER2/ErbB2 (Tyr1248) (1:1000) (#2247, Cell Signaling Technology, USA), rabbit anti-HER3/ErbB3 (1B2) (1:1000) (#4754, Cell Signaling Technology, USA), rabbit anti-phospho-HER3/ErbB3 (Tyr1289) (D1B5) 1:1000) (#2842, Cell Signaling Technology, USA), rabbit anti-rab11a (1:1000) (#ab65200, Abcam, UK), rabbit anti-rab5 (1:1000) (#ab18211, Abcam, UK), rabbit anti-mTOR (7C10) (1:1000) (#2983T, Cell Signaling Technology, USA), rabbit anti-SNX3 (1:300) (#ab56078, Abcam, UK), goat anti-gapdh (1:1000) (#sc-48167, Santa Cruz Biotechnology, USA), mouse anti-tubulin (1:4000) (#T6074, Sigma-Aldrich, USA). After three final washes in 0.1 % PBS-Tween 20, the membranes were incubated with secondary antibodies: IRDye® 800CW Donkey anti-Rabbit IgG (H + L), IRDye® 680RD Donkey anti-Goat IgG (H + L), and IRDye® 680RD Donkey anti-Mouse IgG (H + L) from Li-Cor Biosciences(USA), diluted at 1:10000. Membranes were then developed using the ChemiDoc imaging system from Biorad(USA). Intensities of the bands were then analyzed using the ImageJ gel analyzer tool (ImageJ Software, National Institute of Health, Bethesda, MD, USA, http://rsb.info.nih.gov/ij). A plot profile of western blots was established, and the intensities of all bands were measured. For each sample, the peak intensity was calculated for each target protein by dividing the target protein intensity with the intensity of the loading control protein. Data were then normalized to the control sample.

### Immunoprecipitation

Co-immunoprecipitation (co-IP) experiments were performed using the Dynabeads Protein G Immunoprecipitation Kit (#10007D, Thermofisher Scientific, USA) following the manufacturer’s protocol. Briefly, HEK293 cells overexpressing His-V5-tagged human FRABIN were washed with PBS 72 h post-transfection (with plasmid pEx-FGD4-His-V5), and then incubated in a home-made lysis buffer (HEPES 50 mM, NaCl 150 mM, MgCl2 1.5 mM, EGTA 1 mM, Glycerol 10 %, Triton X100 0.1 %, protease inhibitor). First, 10ug of primary mouse anti-V5 antibody (#ab27671, Abcam, UK) diluted in PBS-Tween 0.01% were incubated with magnetic beads for 1h. The antibody-magnetic beads complex was then incubated with the lysate containing 1 mg of proteins, overnight with end over end rotation. The flow-through is then discarded after placing the tube containing the beads-antigen-antibody lysate, on the magnet. Bound proteins were then eluted with elution buffer and dissociated in 10 µl of NuPAGE LDS Sample Buffer (#NP0007, ThermoFisher Scientific, USA). Samples were then boiled at 70 °C for 10 min before loading on a NuPage Bis-Tris gel.

### Yeast two-hybrid (Y2H)

GAL4-based yeast two-hybrid assay was performed, using a commercial human fetal brain cDNA library containing cDNAs fused to the gal4 activation domain of pEXP-AD502 (ProQuest™, ThermoFisher Scientific, USA), as prey, and full length human *FGD4* cDNA as a bait. To this purpose, the full-length coding sequence of *FGD4* (NM_139241) was subcloned from the pENTR-FL-hFGD4, into a pDBA vector, using the Gateway technology (Thermofisher Scientific, USA). The bait plasmid was transformed in MAV03 yeast strain (MATα; leu2-3,112; trp1-901; his3Δ200; ade2-101; gal4 Δ; gal80 Δ; SPAL10UASGAL1::URA3, GAL1::lacZ, GAL1::His3@LYS2, can1R, cyh2R) following the previously described transformation protocol (Walhout and Vidal 2001). This bait did not show self-activation and was further used for screening. MAV203 cells were then transformed with the prey cDNA library as described (Walhout and Vidal 2001). Following transformation with the cDNA library, yeasts were plated onto synthetic complete (SC) medium minus leucine (−L), minus tryptophane (−W), minus histidine (−H) +25 mM 3-amino-1,2,4-triazole (3-AT) and, incubated at 30°C for 4–5 days. Positive clones were patched onto SC-WHL + 3-AT in 96-well plates, incubated for 3 days at 30°C and transferred in liquid SC-WL for 3 days at 30°C with agitation to normalize the yeast cell concentration used for the phenotypic assay. Cells were then diluted 1/20 in water, spotted onto a selective medium (−WHL+25 mM 3-AT or -WUL), and incubated at 30°C for 4 to 5 days. To perform the β-galactosidase assay, undiluted yeast cells were spotted onto YPD (yeast extract peptone dextrose) medium plates with nitrocellulose filters, and β-galactosidase activity was evaluated one day after. Positive clones were sequenced by Sanger sequencing, after PCR amplification using the following primers: forward: 5’-CGCGTTTGGAATCACTACAGGG-3’ and reverse: 5’-GGAGACTTGACCAAACCTCTGGCG-3’). The clones were identified by using BLAST.

### Statistics

The applied statistical tests, as well as the number of replicates, are indicated in the figure legends. For all experiments based on primary co-cultures, results were obtained from at least three independent cell cultures.

Animals were matched by gender, genotype, and age. Mice were randomly assigned to experimental groups, and gait experiments, as well as processing and analysis of the tissues, were performed in blind. The significance of the results for electron microscopy, primary cultures and in *vivo* niacin treatment was evaluated with one-way or two-way ANOVA or two-tailed student’s t-test according to experimental design. Statistical analyses were performed using GraphPad Prism. All data are presented as mean ± standard error of the mean (SEM).

For gait analysis, each mouse was characterized by the median score of the left and right paw for motor function studies. The scores were analyzed according to the ANCOVA statistics with weight at the corresponding age as a covariate, using (cohen 1988) and SPSS Statistics 19(Field 2005). We calculated the effect size and its confidence interval. Results were expressed as means and SEM.

## Results

### Specific loss of FRABIN in Schwann cell induces myelin abnormalities in vitro and in vivo

To study the role of *FGD4*/FRABIN in CMT4H pathology, we generated a conditional null allele mouse model (**Figure 1A**). To this aim, exon 4 of *Fgd4* was flanked with LoxP sites by homologous recombination. Since previous work from Horn et al (Horn, Baumann et al. 2012) show that the clinical phenotype and the molecular basis of CMT4H relied on the loss of function of FGD4/FRABIN in SCs, without detectable primary contributions from neurons, we abolished FRABIN specifically in Schwann Cells: the conditional mouse model is referred to as *Fgd4*^*SC-/-*^. Conditional ablation of *Fgd4* in SCs was obtained by breeding mice homozygous for the floxed allele (*Fgd4*^*flox/flox*^) with transgenic P0-Cre mice to drive expression of the Cre-recombinase in SCs at embryonic day 13.5 (E13.5) (Feltri, D’Antonio et al. 1999). We demonstrated the effective deletion of the floxed exon 4 in this conditional knock-out *Fgd4*^*SC-/-*^ mouse model, by RT-PCR using total RNA extracted from sciatic nerves of *Fgd4*^*SC-/-*^ and control mice (**Figure 1B-C**). As expected, we show that the deletion of exon 4 leads to a shorter fragment amplified from between exons 3 and 7 in the mutant nerves (374 bp in *Fgd4*^*SC-/-*^ versus 882 bp in the WT). Due to the lack of antibodies recognizing specifically mouse FRABIN, we were, unfortunately, unable to assess the deletion by Western-Blot.

**Figure 1.**
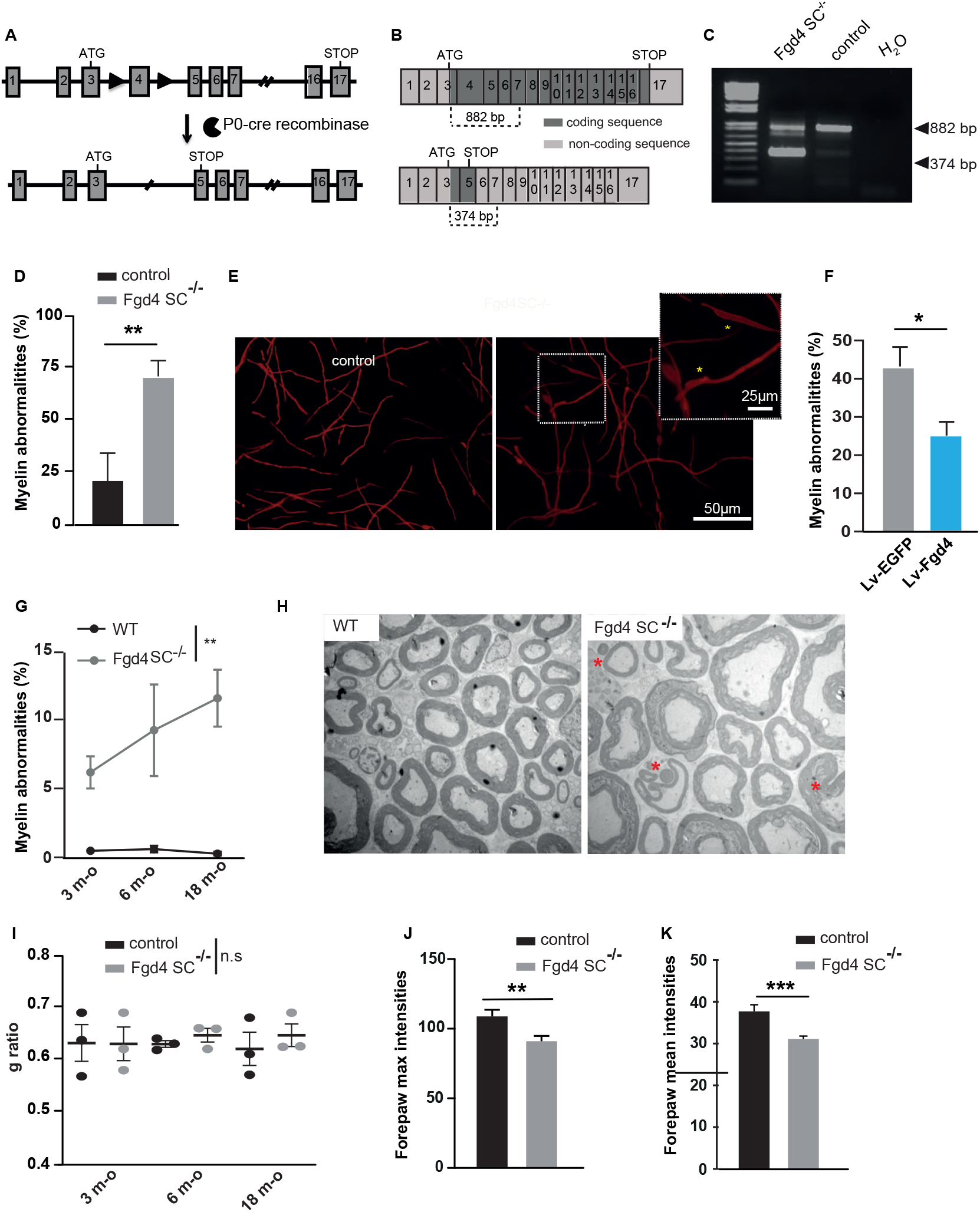
Generation and phenotypic characterization of a new mouse model of CMT4H. (**A-C**) Generation of a conditional knock-out mouse model for CMT4H. (**A**) The floxed allele (Fgd4^fl/+^) was generated by flanking exon 4 with loxP sites (black arrowheads) in the *Fgd4* gene leading to a frameshift by generation of a premature stop codon in exon 5. The conditional deletion of *Fgd4* in SCs is induced by crossing *Fgd4*^*fl/fl*^ mice with mice expressing the cre recombinase under the Mpz/P0 promoter. For simplification the conditional knock-out *Fgd4*^*fl/fl*^ ; P0cre mice are referred to *Fgd4*^*SC-/-*^. (**B**) *Fgd4* transcript before (up) or after (bottom) excision of exon 4 following cre-recombinase expression. (**C**) Excision of the exon 4 from *Fgd4* transcript, was verified, by RT-PCR, using RNA extracted from the sciatic nerves of *Fgd4*^*SC-/-*^ and control mice. Due to exon 4 excision, the size of the amplified amplicon was 882 bp and 374 bp in WT and *Fgd4*^*SC-/-*^ sciatic nerves respectively. (**D-E**) Loss of Fgd4/FRABIN alters myelination *in vitro*. **(D)** Number of myelin anomalies quantified in control and *Fgd4*^*SC-/-*^ cocultures. The presence of focal hypermyelination defects was evaluated in 150-160 myelinated segments per coverslips and 3 independent cultures. Myelinated segments were visualized by immunolabeling of Myelin Basic Protein (MBP). Data are expressed as mean ± sem. Statistical analysis: two-way ANOVA with Sidak post-hoc test. (**E**) Illustration of myelinated segments in control and *Fgd4*^*SC-/-*^ cocultures. Examples of myelin abnormalities that can be observed in the enlarged image and identified by asterisks (*). **(F)** Overexpression of Fgd4/FRABIN in *Fgd4*^*SC-/-*^ cocultures reduces the proportion of abnormally myelinated fibers. *Fgd4*^*SC-/-*^ cocultures were infected with Lv-EGFP or Lv-Fgd4, 1 day after seeding. Data are expressed as mean percentage ± sem (n= 4 independent cultures). Statistical analysis: unpaired Student’s t-test. (**G-H**) Loss of Fgd4/FRABIN alters PNS myelination *in vivo*. (**F**) Proportion of abnormal myelinated fibers in the distal part of sciatic nerve of 3, 6 and 18 months-old WT and *Fgd4*^*SC-/-*^ mice (n= 3 animals per genotype). Data are expressed as mean percentage ± sem. Statistical analysis: two-way repeated-measures ANOVA (group*time) with Sidak post-hoc test (**H**) Electron microscopy pictures illustrating myelination of sciatic nerves of WT (left panel) and *Fgd4*^*SC-/-*^ (right panel) mouse. Asterisks indicate outfoldings. scale bar : 5 µm. (**I**) g-ratio analysis revealed no statistical difference in myelin thickness in the sciatic nerve of at 3, 6 and 18 mo WT and *Fgd4*^*SC-/-*^ mice. A total of 500-1000 axons between 0.5 and 6 µm for animals (n=3 per genotype) were analysed. Data are presented as mean ± sem (n=3 animals). (**J**-**K)** Max and mean forepaw intensities are significantly decreased in Fgd4^SC-/-^ mice compared to WT mice. Data are expressed as mean ± sem (n=11 WT and n=13 Fgd4^SC-/-^ mice). * p<0.05, ** p<0.01, ***p<0.001.

We previously observed that the loss of FRABIN leads to aberrant myelin structures called outfoldings in nerves of patients affected with CMT4H (De Sandre-Giovannoli, Delague et al. 2005, Delague, Jacquier et al. 2007, Boubaker, Hsairi-Guidara et al. 2013). Here, we wanted to assess whether such abnormalities could be reproduced in our *Fgd4*^*SC-/-*^ mouse model both *in vitro* and *in vivo. In vitro*, we established Dorsal root ganglia (DRG)/SC cocultures from E13.5 *Fgd4*^SC-/-^ embryos, in which myelination was induced for 15 days using ascorbic acid. Myelinated segments were visualized by Myelin Basic Protein (MBP) immunostaining (**Figure 1D-E**). We thus noticed irregularly shaped MBP-positive fibers reflecting focal hypermyelination (**Figure 1E**), similar to the ones previously described in a coculture model derived from the mouse model for CMT4B1 (Bolis, Coviello et al. 2005, Bolino, Piguet et al. 2016). In *Fgd4*^*SC-/-*^ cocultures, the proportion of these abnormal MBP+ fibers represented 71.4 ± 4.3% of total MBP+ fibers, as compared to control cocultures, where they represent 21.6 ± 8.3% (**Figure 1D-E**). To determine whether the presence of those myelin defects was related to a loss of *Fgd4*/FRABIN, we overexpressed *Fgd4/FRABIN* in our mutant *Fgd4*^*SC-/-*^ coculture model using lentivirus. Overexpressing *Fgd4/FRABIN* led to a significant decrease in the proportion of abnormal MBP+ segments in mutant *Fgd4*^*SC-/-*^ cocultures (39.8 ± 5.6%) to a level comparable to control conditions (23.1 ± 2.9%) (**Figure 1F)**. This finding confirms that loss of *Fgd4*/FRABIN in SCs is a major contributor to the altered myelin phenotype.

Altered myelination was in parallel evaluated, over-time, *in vivo*, in the sciatic nerves of both *F*gd4^SC-/-^ and control mice. *In vivo*, we observed out and infoldings as described in the nerves of CMT4H patients. We observed a higher proportion of abnormal myelinated fibers in *F*gd4^SC-/-^ ‘s nerves as compared to control, as soon as 3 month-old, which increased significantly over time (3 month-old: control 0 ± 0,1% and *F*gd4^SC-/-^ 4.9 ± 1.9%; 6 month-old: control 0.46 ± 0.32% and Fgd4^SC-/-^ 9.1 ± 5.1%; 18 month-old: control 0.44 ± 0.26% and *F*gd4^SC-/-^ 11.7 ± 2.1 %) (**Figure 1G-H**). The presence of altered myelin was not correlated to a defect of myelin thickness as the g-ratio was not affected at those studied time points (**Figure 1I and Figure Sup1A)**. Finally, in addition to myelination defects, 12 months-old *Fgd4*^*SC-/-*^ mice display significant denervation of the neuromuscular junctions (NMJs) of the gastrocnemius muscle (% of fully innervated NMJs: WT 75,3 ± 3,1% versus *Fgd4*^*SC-/-*^ 50,3± 7%; % of intermediate NMJs: WT 14,5 ± 1,9% versus Fgd4^SC-/-^ 27,06± 6,6% and % of denervated NMJs: WT 10,1 ± 1,4% versus Fgd4^SC-/-^ 22,6± 1,6%;), suggesting late distal axonal degeneration (**Figure sup 1B-C**).

To assess the locomotor performance of our CMT4H model, we monitored footprint analyses using the gait test at 6, 12 and, 18 month-old. Impaired motor functions were detected in the oldest mice only, as evidenced by a difference in parameters reflecting the pressure exerted by each paw (i.e intensity parameters). Indeed, the Maximal forepaw intensity was lower in F*gd4*^*SC-/-*^ mice: 92.4 ± 3.9 compared to controls: 110.6 ± 4.8 (F = 8.37, df 1, 24, p < 0.08) as well as the mean forepaw intensity: F*gd4*^*SC-/-*^: 31.08 ± 0.7 vs WT 37.7 ± 1.6 (F = 17.28, df 1, 24, p < 0.001). These differences exceed the limits between pathology and typicality as shown by the large effect’s sizes (*η*^*2*^ = 0.28, CI from 0.11 to 0.57 and *η*^*2*^ = 0.43, CI from 0.19 to 0.53, respectively) (**Figure 1 J-K**).

### Specific loss of FRABIN in Schwann cell leads to a deregulation of the Neuregulin1 type III-Erbb2/3 myelination pathway

Previous studies reported that myelin outfoldings might arise from an enhanced NRG1 type III/ErbB2/3/ AKT/mTOR pathway activation (Goebbels, Oltrogge et al. 2012, Bolino, Piguet et al. 2016). We, therefore, assessed by western blot, the level of expression of key components of this pathway, both *in vitro* and *in vivo* (**Figure 2**). *In vitro*, we noticed a significant increase in the levels of phosphorylated Erbb2 receptor (P-Erbb2), phosphorylated AKT (P-Akt) and mTOR, in *F*gd4^*SC-/-*^ cocultures as compared to control (**Figure 2A-B)**. The level of full-length and cleaved forms of NRG-1 type III was not affected (**Figure Sup 1D-E**). Abnormal regulation of the NRG1 type III-ErbB2/3 pathway was confirmed *in vivo*, in the sciatic nerves of full knock-out mice (F*gd4*^*-/-*^). We used full knock-out animals (F*gd4*^*-/-*^), rather than conditional knock-out animals (F*gd4*^*SC-/-*^) because, in the latter, the normal expression of Frabin in the axons (which are not knock-out) may compensate for any change in the level of expression of the NRG1 type III/ErbB2/3 pathway due to the knock-out in Schwann cells. In F*gd4*^*-/-*^ sciatic nerves, we observed, by western-blot, a significant upregulation of Erbb2 and mTOR levels, as compared to control nerves, while levels of AKT and its phosphorylated form were not different. We also noticed a significant decrease of the full-length NRG1 type III (i.e., the pro-protein) and a concomitant increase of the levels of the C-terminal NRG1 fragment (Falls 2003), which indicates an increase in the cleavage of the NRG1 type III pro-protein. Overall those observations demonstrate an enhanced activation of the NRG1 type III/ErbB2/3/ AKT/mTOR pathway (**Figure 2C-D and Figure sup 1 F-G**).

**Figure 2.**
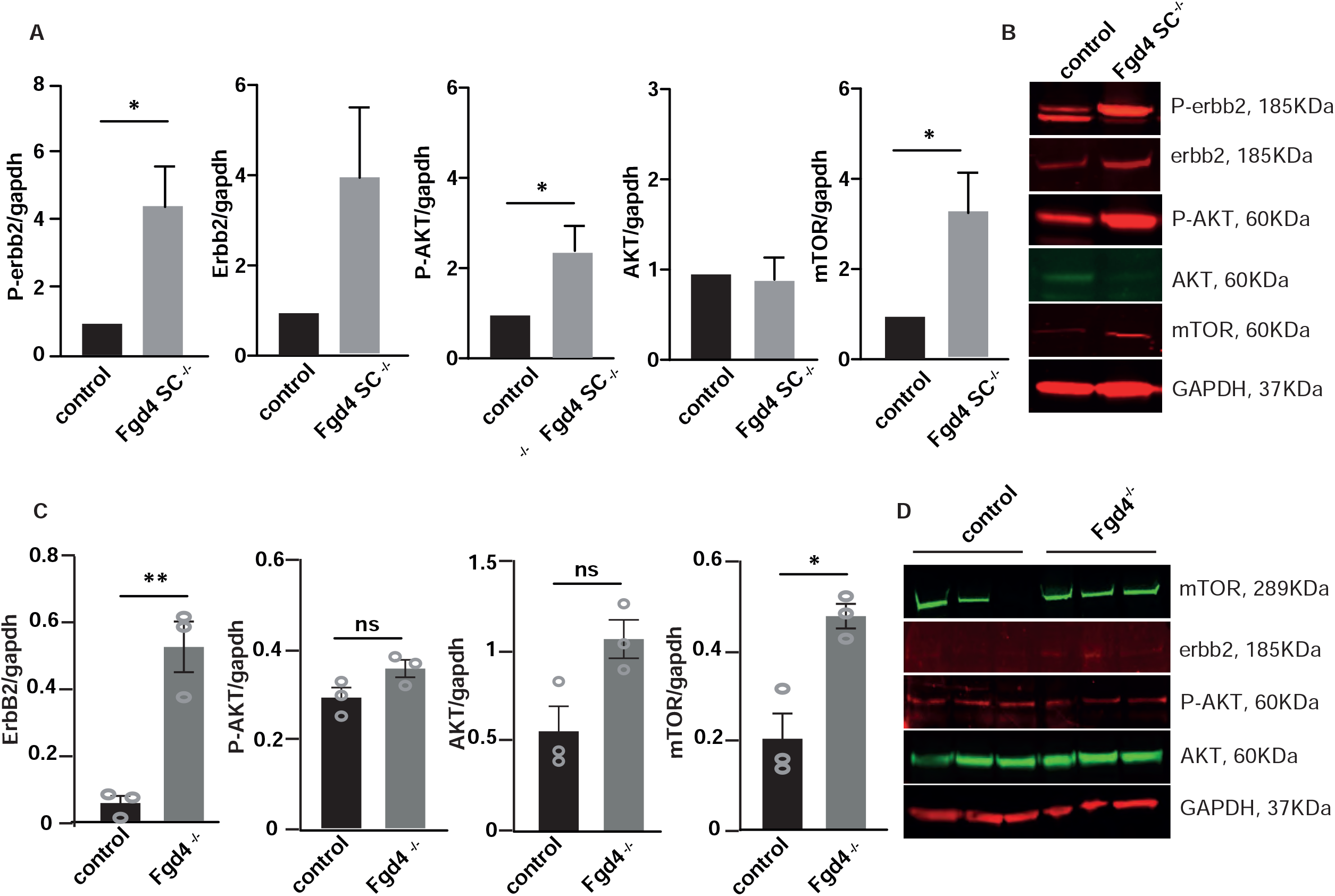
Neuregulin-1 type III/ErbB2/3/AKT/mTOR pathway is upregulated in CMT4H models. (**A-B**) Phosphorylated ErbB2 (P-ErbB2) and AKT (P-AKT) as well as mTOR expression are significantly increased in Fgd4^SC-/-^ cocultures compared to control. (**A**) Levels of expression of P-ErbB2, ErbB2, P-AKT, AKT and mTOR were assessed by western-blot analysis. Data are expressed as mean ± sem (n=3-4 cocultures). Statistical analysis: unpaired Student’s t-test. (**B**) Western blot pictures illustrating the expression of the markers described in (A). (**C-D**) Levels of expression of P-ErbB2, ErbB2, P-AKT, AKT and mTOR were assessed by western-blot analysis in the sciatic nerves of Fgd4^-/-^ mice compared to WT mice. (**C**) Data are expressed as mean ± sem (n=3 animals per genotype). Statistical analysis: unpaired Student’s t-test. (**D**) Western blot pictures illustrating the expression of the markers described in (E). * p<0.05, ** p<0.01, ***p<0.001.

### Loss of FRABIN do not affect the myelination-related gene expression program

To examine global gene expression involved in the myelination abnormalities linked to the absence of FRABIN in SCs, we performed bulk RNA-Sequencing on duplicates from *Fgd4*^*SC-/-*^ and control WT cocultures. Heatmap of unsupervised hierarchical clustering of the 4 samples (2 *Fgd4*^*SC-/-*^ and 2 WT) showed that *Fgd4*^*SC-/-*^ samples clustered separately from the controls (**Figure 3A**). Gene Set Enrichment Analysis (GSEA) on the entire set of expression data, highlighted a total of 2451 genes differentially expressed genes (DEG) between *Fgd4*^*SC-/-*^ and control cocultures. Among them, 1679 were overexpressed and 772 underexpressed (**Figure 3B**).

**Figure 3.**
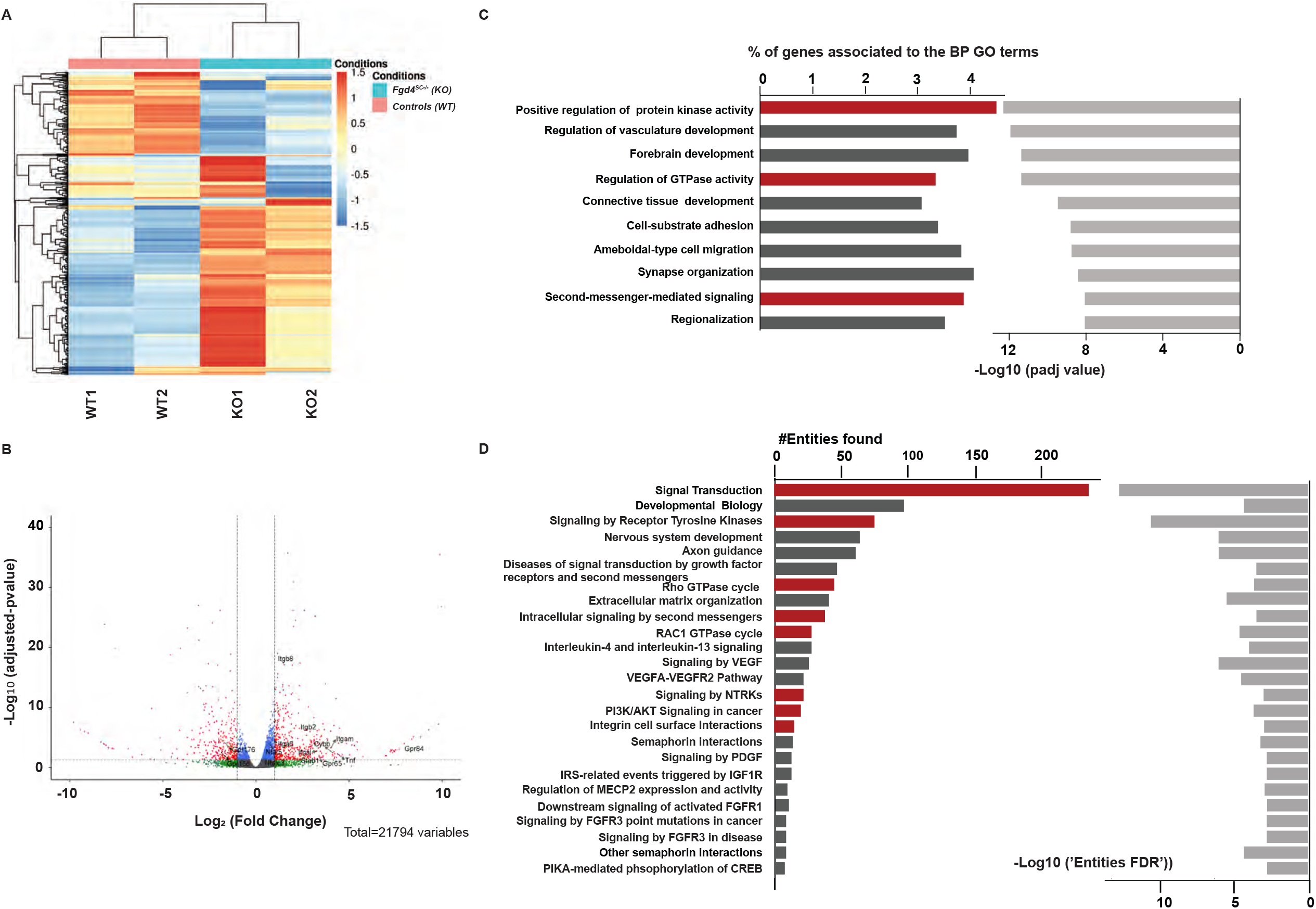
Transcriptional profiles of *in vitro* myelin samples (DRG/SC cocultures) from conditional Knock-out (*Fgd4*^*SC-/-*^) and control (WT) mice. (**A**) Full heat map of unsupervised hierarchical clustering of the 4 samples (n= 2 replicates for *Fgd4*^*SC-/-*^ and 2 replicates for WT). The scale bar unit is obtained applying a variance stabilizing transformation (VST) to the count data (DESeq2:: VarianceStabilizingTransformation) before normalization. (**B**) Volcano plots showing the distribution of gene expression fold changes and adjusted p-values between the two conditions. A total number of 21,794 genes were tested. Padj < 0.05 was used as the threshold to reject the null hypothesis and consider the difference in gene expression. Red plots represent genes significantly deregulated genes (adjusted p-value<0.05, with fold change (FC) >2 or <-2). Genes with significant deregulation (adjusted p-value<0.05) but with small FC (−2<FC<2) are indicated in blue. Green and gray dots represent genes with non-significant Fold Changes (adjusted p>0.05).(**C**) Top 10 Biological Process (BP) Gene Ontology (GO) terms enriched in the DEGs. We identified biological pathways (BPs) with an adjusted p-value lower than 0.05. The bars on the left represent the percentage of DEGs determined for each represented GO term from the total number of DEGs. Light gray bars on the right represent the enrichment score (−Log10 of adjusted p-value) for each GO term. GO terms with particularly interesting functions, regarding the NRG1 type III pathway and FRABIN are highlighted in red. (**D**) Top 25 reactome pathways enriched for the 536 genes found in the top 10 significantly enriched GO terms from **C**), (adj. p-value < 0.001). The bars represent the number of DEGs found in each reactome pathway. Pathways with particularly interesting functions, regarding the NRG1 type III pathway and FRABIN are highlighted in red.

Functional Gene ontology (GO) annotation of the DEGs, with statistical significance of padj value < 0.05, showed enrichment in several Biological Processes (BP). Interestingly, in the top enriched BP GO terms, we found “positive regulation of protein kinase activity” (GO:0045860), “regulation of GTPase activity” (GO:0043087), “second-messenger-mediated signaling” (GO:0019932), or “cell-substrate adhesion” (**Figure 3C**), which are related to important processes in myelination or to the known functions of FRABIN (regulation of GTPase, binding to PIs). Enrichment analysis of the Reactome Pathways using the list of the 536 DEGs from the Top10 enriched GO terms indeed confirmed major deregulations in signal transduction in *Fgd4*^*SC-/-*^ coculture. As expected, considering the known function of FRABIN as a GEF for small RhoGTPases, we observed enrichment of genes encoding proteins involved in the RAC1 GTPase cycle (#R-HSA-9013149), Rho GTPase cycle (#R-HSA-9012999), and Intracellular signaling by second messengers (#R-HSA-9006925)(**Figure 3D**). In addition this analysis of pathway enrichment highlights new signaling transduction pathways, which might be important for correct myelination, such as Integrin cell surface interactions (**Figure 3D**).

In-depth analysis of DEGs from the top enriched GO terms revealed several deregulated genes, such as members of the integrin family (in particular Itgb8), members of the G protein-coupled receptor (Gpr) family, such as Gpr65 and Gpr84, or other signaling proteins (Nfat5, Tnf, Cybb, stab1). These genes are highlighted in **Figure 3B**. By looking more closely at the DEGs, we did not reveal any changes of expression in NRG1 type III/ErbB2/3 pathway and associated direct downstream factors, nor in myelin structural genes.

### Specific loss of FRABIN in Schwann cell impairs endosomal trafficking

Emerging data propose that dysregulation of ErbB receptor trafficking may be the cause of the impairment of NRG type III/ErbB2/3 signaling in SCs and represent a common pathogenic mechanism in several CMT4 subtypes (Lee, Chin et al. 2017). FRABIN is a GEF responsible for GDP/GTP exchange on the small RhoGTPases cdc42 and Rac1, and also contains two Pleckstrin Homology and one FYVE domains, that are implicated in interactions with phosphoinostides (PIs). PIs are lipid regulators of endocytic trafficking (Cullen and Carlton 2012). Previous work by Horn et al (Horn, Baumann et al. 2012) has shown an impairment of endocytosis in the absence of FRABIN, but its impact on myelination defects has not been studied.

To determine whether the myelination defects observed in the *F*gd4^SC-/-^ mice are connected to defective endocytic trafficking, we looked at the expression of the early endosomal marker RAB5 and the recycling endosomal marker RAB11, *in vitro*. In F*gd4*^*SC-/-*^ cocultures, RAB5 expression levels are increased, as compared to controls but without reaching statistical significance. In contrast, RAB11 is significantly upregulated in conditions of *F*gd4^SC-/-^ cocultures compared to control (**Figure 4A-B**). To follow more precisely the endocytic trafficking following FRABIN’s loss, we monitored the trafficking of pHrodo-Transferrin (phrodo-TF), over time, in primary SCs knocked-down (KD) for *Fgd4/FRABIN*, as well as in control cells. We induced *Fgd4/FRABIN* KD using a shRNA targeting*Fgd4* (SHFgd4), previously described by Horn et al. (Horn, Baumann et al. 2012), that we delivered in rat primary SCs using a lentivirus (**Figure sup2C**). SCs were then pulse-labeled, 48h post-infection, with fluorescent pHrodo-TF for 30 min at 4°C to prevent uptake and then chased by incubating at 37°C to allow uptake (15 min) and recycling (30 and 45 min). pHrodo-TF has the property to become fluorescent once internalized in the endosomes. By quantifying the levels of fluorescence of pHrodo-TF, we noticed similar levels of internalized TF in SCs expressing a control shRNA (SHcontrol) or SHFgd4, after 15 min of incubation, suggesting a similar capacity of TF-endocytosis between the control and Fgd4 KD conditions (**Figure 4C-D**). In both control and Fgd4 KD conditions, the level of fluorescence of phrodo-TF decreases over time due to TF recycling back to the membrane. Interestingly, in comparison to the control conditions, the levels of fluorescence of pHrodo-TF are significantly lower after 30 and 45 min of incubation in Fgd4 KD conditions (**Figure 4C-D**). This decrease in TF-retention in the endosomes suggested an increase of pHrodo-TF recycling back to the membrane in conditions of loss of *Fgd4*/FRABIN.

**Figure 4.**
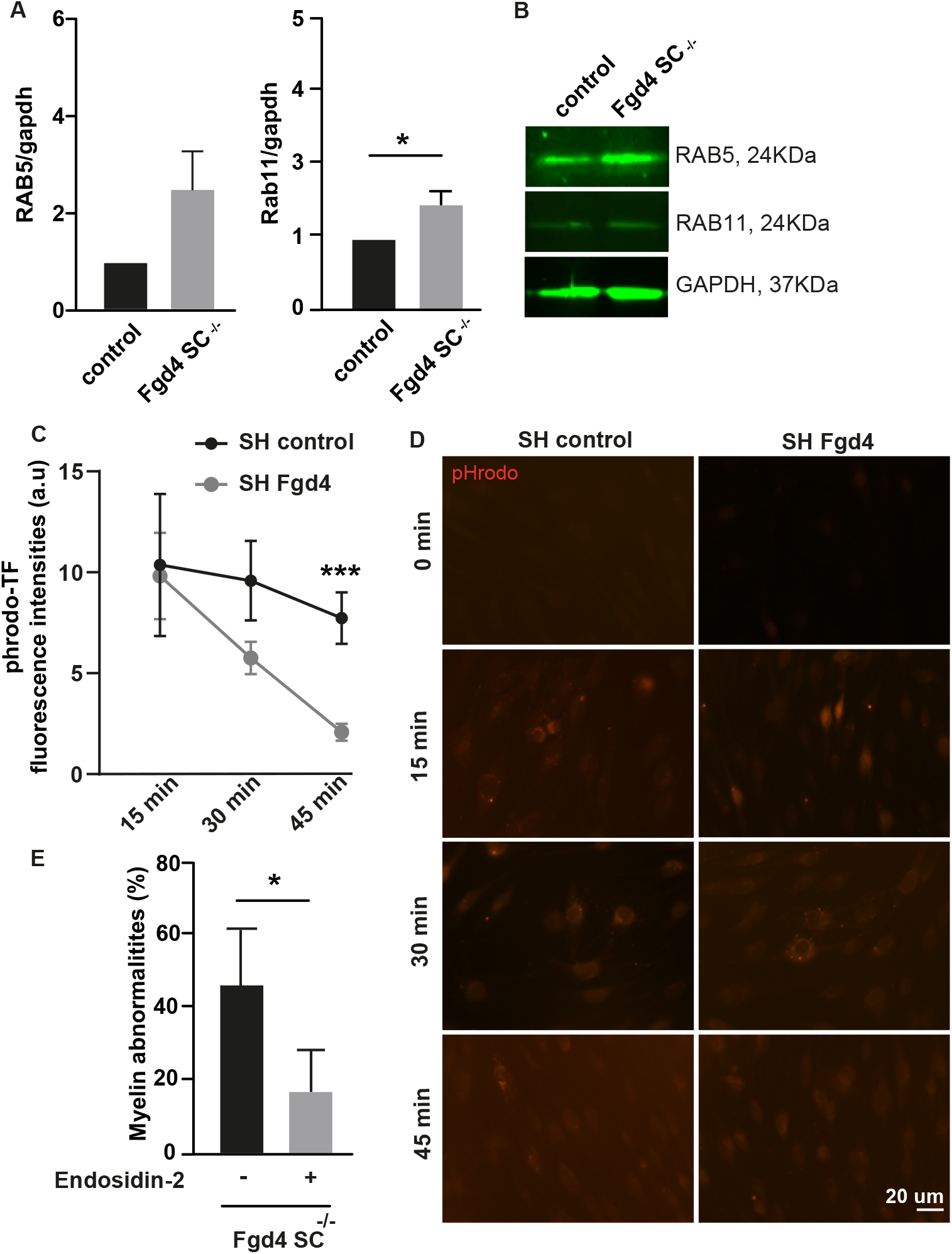
Altered endocytic trafficking contribute to the abnormal myelination observed in *Fgd4*^*SC-/-*^ conditions. (**A-B**) The expression of two endosomal markers (i.e RAB5: early endosome and RAB11: recycling endosome) is upregulated in *Fgd4*^*SC-/-*^ cocultures, as compared to control. (**A**) Quantification of RAB5 and RAB11 expression levels in Western Blot observed in (B). Data are expressed as mean ± sem (n=3 cocultures). Statistical analysis: unpaired Student’s t-test. (**B**) Illustration of RAB5 and RAB11 expression patterns observed by western-blot in control and *Fgd4*^*SC-/-*^ cocultures. (**C-D**) pHrodo-Transferrin (pHrodo-TF) trafficking is altered in primary SCs infected with a Lv expressing a shRNA control or directed against *Fgd4*. 48h after Lv infection, SCs were incubated with pHrodo-TF during 15, 30 or 45 min and immediately fixed in PFA 4%. pHrodo-TF is weakly fluorescent at a neutral pH, but brightly fluorescent in acidic compartments such as endosomes. (**C**) pHrodo-TF fluorescence was quantified in infected cells (n=60-80 cells for each condition, from 3 independent-cultures) and data are expressed as mean ± sem for each time point. Statistical analysis: unpaired Student’s t-test for each time-points. (**D**) Examples of pHrodo-TF labeling after 0, 15, 30 or 45 min in control (shRNA-control) and knocked-down (shRNA-Fgd4) conditions. (**E**) The blocking of endosomal recycling using Endosidin-2 (1µM) reduces significantly the proportion of abnormal myelin in *Fgd4*^*SC-/-*^ cocultures compared to non-treated conditions. Data are expressed as mean percentage ± sem (n= 3 cocultures). Statistical analysis: unpaired Student’s t-test. * p<0.05, ** p<0.01, ***p<0.001.

To assess whether the observed endocytic trafficking defect could participate in the improper myelination observed in the *F*gd4^SC-/-^ cocultures, we treated the cocultures with endosidin-2 (ES-2, 1µM), a compound known to reduce exocytosis and endosomal recycling (Zhang, Brown et al. 2016). Interestingly, treating F*gd4*^*SC-/-*^ cocultures with ES-2 decreases significantly the number of myelin abnormalities observed in conditions of loss of FRABIN (non-treated-*F*gd4^SC-/-^: 47 ± 6.8%, ES-2 treated Fgd4^SC-/-^: 17.5 ± 5.7%) (**Figure 4E**). Altogether, these data support a defect in endosomal trafficking following Fgd4/FRABIN loss which may compromise proper myelination signaling and contribute to the abnormal myelination observed *in vitro*.

### Sorting nexin 3, a new partner of FRABIN, contributes to CMT4H pathogenesis

As FRABIN’s function in vesicle trafficking remains poorly studied, we searched for FRABIN interactors that could be involved in this process. To achieve this, we performed a yeast two-hybrid screening analysis in a human fetal brain library, using Fgd4 cDNA as bait. Among the partners identified, we focused on Sorting nexin 3 (SNX-3). SNX-3 belongs to the Sorting nexin protein family implicated in membrane trafficking. SNX3 is associated with the early endosome through its PX domain and interacts, as FRABIN, with Polyphosphoinositide PI3P (Xu, Hortsman et al. 2001). In addition, SNX-3 is known to facilitate the recycling of the Transferrin receptor (Chen, Garcia-Santos et al. 2013). We first assessed the localization of FRABIN and SNX3 in the S16 rat SC line by immunofluorescence. We noticed vesicular staining of the endosomal marker SNX3, as previously described by (Xu, Hortsman et al. 2001), and colocalization with FRABIN, especially at the perinuclear region (**Figure 5A**). In parallel, we assessed their potential interaction by co-immunoprecipitation. HEK293 cells previously transfected with FRABIN-His-V5 were subjected to immunoprecipitation using an anti-V5 antibody. Western blot analysis revealed that SNX3 was co-immunoprecipitated in the lysate fraction (**Figure 5B)**.

**Figure 5.**
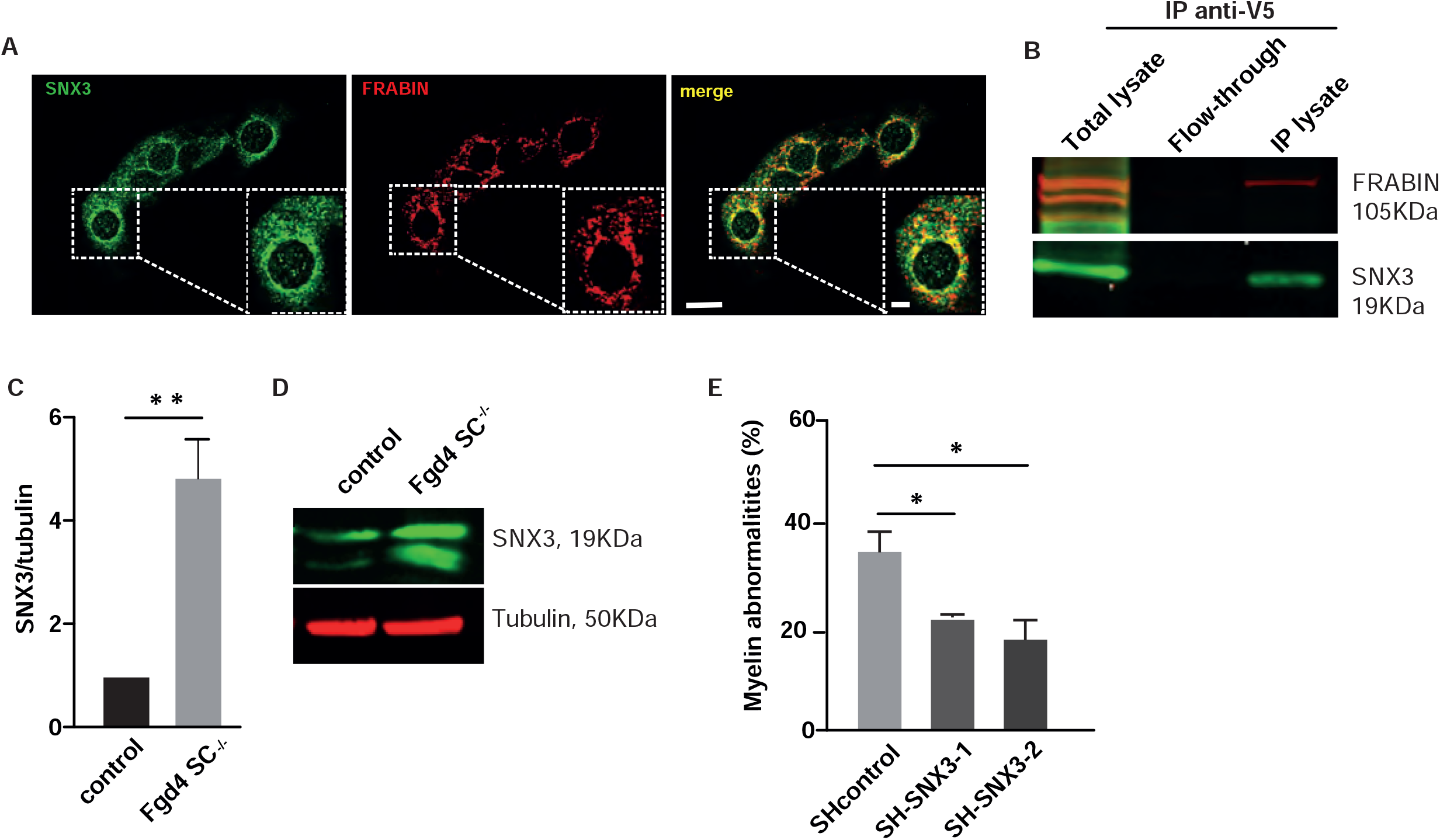
SNX3, a new partner of FRABIN, contribute to abnormal myelination in vitro. (**A**) FRABIN (red) colocalizes with the endosomal marker SNX3 (green) in the immortalized S16 rat SC line. Colocalization areas are visualized in yellow in the merge picture. Scale bar of the whole picture : 20 µm, scale bar of the enlarged picture : 10 µm. (**B**) Co-immunoprecipitation (IP) of SNX3 and FRABIN in HEK293 cells overexpressing V5-tagged FRABIN.SNX3 was detected using an anti-SNX3 antibody after immunoprecipitation of V5-FRABIN using an anti-V5 antibody. Note that both FRABIN and SNX3 are found only in the IP lysate, and not in the flow-through fraction. (**C-D**) SNX3 levels are increased in *Fgd4*^*SC-/-*^cocultures. SNX3 levels of expression were evaluated by western-blot in both control and *Fgd4*^*SC-/-*^ cocultures. (**C**) Quantification of proteins levels of SNX3 observed by Western Blot. Data are expressed as mean ± sem (n=3 cocultures). Statistical analysis: unpaired Student’s t-test. (**D**) Western-blot illustration of SNX3 expression profile, normalized to Tubulin. (**E**) Knock-down of *Snx3* reduces significantly the number of myelin abnormalities in *Fgd4*^*SC-/-*^ cocultures. *Fgd4*^*SC-/-*^ cocultures were infected at day 1 after plating with Lv expressing either SH control, or SH-SNX3 (1 and 2). Myelination was then induced by ascorbic acid (50µM) treatment during 12 days. Data are expressed as mean ± sem (n=3 cocultures). Statistical analysis: one-way ANOVA, with Sidak post-hoc test (multiple comparison to SH control).

We then examined the levels of SNX-3 in control and *F*gd4^SC-/-^ cocultures by western-blot. We observed a significant increase in the level of expression of SNX-3 in *F*gd4^SC-/-^ cocultures compared to control (**Figure 5C-D**). To determine whether the upregulation of SNX3 was contributing to CMT4H pathology, we downregulated the expression of SNX3 using RNA interference in *F*gd4^SC-/-^ cocultures. F*gd4*^*SC-/-*^ cocultures were infected with LV expressing either a control shRNA or two different shRNAs targeting *Snx3*. The efficiency of SNX3 knock-down was evaluated one-week post-infection by western-blot (**Figure sup 2A-B**). By quantifying the number of MBP+ segments harboring myelination abnormalities, we noticed a significant reduction in their percentage following *Snx3* knock-down in *Fgd4*^*SC-/-*^ cocultures with a KD of *Snx3*: 35.2% ± 4 in *F*gd4^*SC-/-*^ cocultures + Lv-SH control, 21.9% ± 0.6 in *F*gd4^*SC-/-*^ cocultures+ Lv-SH SNX3-1, or 18% ± 3.9 in *F*gd4^*SC-/-*^ cocultures + Lv-SH SNX3-2 (**Figure 5E**). We conclude that SNX3 is a new partner of FRABIN in SCs and that an increase in SNX3 levels is likely to contribute to the focal hypermyelination events in CMT4H.

### Niacin reduces myelin abnormalities in both in vitro and in vivo models of CMT4H

It is known that the amount of axonal NRG1-type III, and its downstream signaling via the activation of the ErbB2/3 receptors, determines the thickness of myelin sheath (Michailov, Sereda et al. 2004, Taveggia, Zanazzi et al. 2005). It is well-known that nerves from patients with CMT4H display abnormalities characteristic of aberrant hypermyelination, such as outfoldings (De Sandre-Giovannoli, Delague et al. 2005, Stendel, Roos et al. 2007, Boubaker, Hsairi-Guidara et al. 2013). Here we showed that this abnormal myelination observed in the absence of FRABIN is due to abnormal regulation of the NRG1 type III/ErbB2/3 pathway (**Figure 2**). We therefore aimed at lowering myelination by reducing the NRG1 type III/ErbB2/3 signaling, using Nicotinic acid (Niacin). Indeed, niacin has been shown to promote the activation of the *α*-secretase Tace, which leads to a shorter cleaved form of NRG1 type III unable to link its ErbB2/3 receptors (Chen, Cui et al. 2009). Therefore, cleavage of NRG1 type III by Tace negatively regulates PNS myelination. We first tested Niacin treatment *in vitro*, by adding 5 mM of Niacin, concomitantly to ascorbic acid, every two days during 10 days, in *F*gd4^SC-/-^ cocultures. We observed that Niacin treatment reduces by ∼ 60 % the number of myelin defects (*F*gd4^*SC-/-*^ : 71.4% ± 4.4 versus *F*gd4^SC-/^ + Niacin: 28.4% ± 4.6) **(Figure 6A-B)**. We then checked, by western blot, whether the benefit of Niacin treatment was due to a downregulation of NRG1 type III/ErbB2/3 signaling. After Niacin treatment, we observed a significant reduction of the levels of expression of P-ErbB2 and ErbB2 (**Figure 6C-D)**.

**Figure 6.**
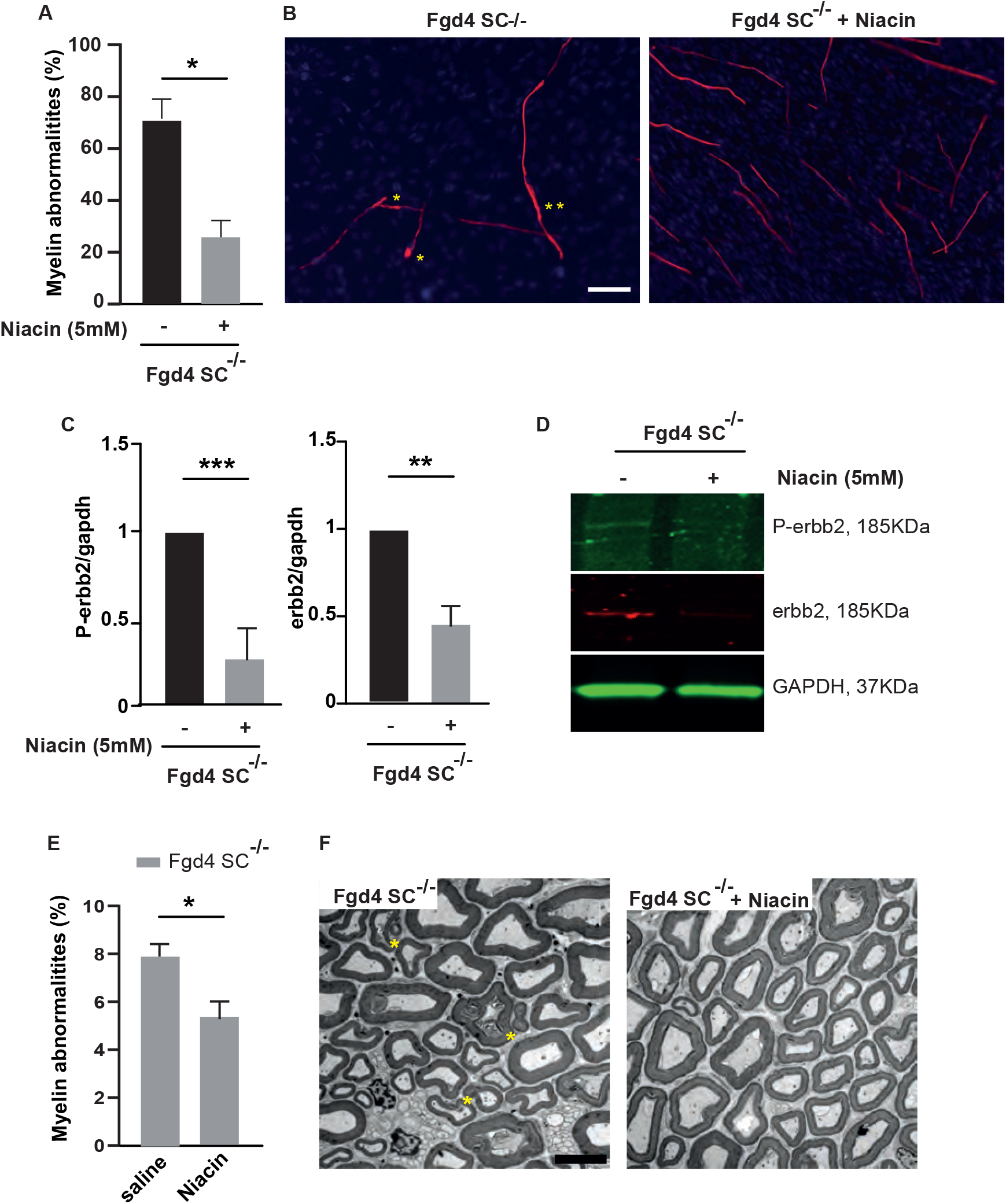
Downregulating NRG1-type III/ErbB2/3 signaling using Niacin reduces myelination defects both *in vitro* and *in vivo*. (**A-B**) Niacin treatment reduces significantly myelin abnormalities in *Fgd4*^*SC-/-*^ cocultures. *Fgd4*^*SC-/-*^ cocultures were treated every two-days with Niacin (5mM) concomitantly to ascorbic acid (50µM) during 12 days. (**A**) Quantification of the myelin abnormalities. Data are expressed as mean ± sem (n=3 cocultures). Statistical analysis: unpaired Student’s t-test. (**B**) Examples of MBP immunostaining in *Fgd4*^*SC-/-*^ non-treated (left panel) and treated with Niacin (right panel) cocultures. Examples of myelin abnormalities are indicated by asterisks (*outfolding, **tomacula). (**C-D**) Niacin treatment leads to a downregulation of the expression of NRG1-type III/ErbB2/3. P-ErbB2, ErbB2, full-lenght and cleaved NRG1 expression were assessed by western-blot in non-treated and niacin-treated *Fgd4*^*SC-/-*^ cocultures. (**C)** Quantification of P-erbb2, erbb2, cNRG1-III and fNRG1-III. Data are expressed as mean ± sem (n=3 cocultures). Statistical analysis: unpaired Student’s t-test. (**D**) Western blot pictures illustrating the expression of the markers described in (C). (**E-F**) Niacin treatment reduces the proportion of abnormal myelin fibers *in vivo*. One month-old *Fgd4*^*SC-/-*^ mice were daily injected with saline solution or Niacin (60mg/Kg) during 8 weeks. Myelin abnormalities were quantified in the distal part of the sciatic nerves of the mice. (**E**) Quantification of the myelin abnormalities in sciatic nerves of treated and non-treated *Fgd4*^*SC-/-*^ mice. Data are expressed as mean ± sem (n=4 saline-treated animals and 5 Niacin-treated animals). Statistical analysis: unpaired Student’s t-test. (**F**) Electron microscopy pictures illustrating myelination of sciatic nerves of saline treated (left panel) and Niacin treated Fgd4^SC-/-^ (right panel) mouse. Asterisks indicate outfoldings. scale bar : 5 µm.

We then evaluated the myelination status of the sciatic nerves of Niacin-treated *F*gd4^SC-/-^ mice compared with saline-injected *F*gd4^SC-/-^ mice. Since we observed a significant proportion of myelin defects as soon as 3 month-old in *F*gd4^SC-/-^mice, mice were treated via IP injections, for 8 weeks starting from 4 week-old (before the onset of myelin defects). Importantly, we noticed a significant reduction of the number of defective myelin fibers in treated versus non-treated animals (*F*gd4^SC-/-^ + Niacin: 5% ± 0.7 versus non-treated *F*gd4^SC-/-^ : 7,4% ± 0.5) **(Figure 6E-F)**.

These observations demonstrate that the overactivation of the NRG1 type III/ErbB2/3 pathway underlies the myelination anomalies linked to CMT4H pathogenesis. Consequently, reducing NRG1 type III/ErbB2/3 NRG1activation via Niacin represents an interesting treatment perspective for CMT4H.

## Discussion

CMT4H is an autosomal recessive demyelinating subtype of Charcot-Marie-Tooth disease, that we have first described in 2005 (De Sandre-Giovannoli, Delague et al. 2005) and for which we and others have described *FGD4* as the gene causing the disease (Delague, Jacquier et al. 2007, Stendel, Roos et al. 2007). One distinctive feature of CMT4H is the presence, in the nerves of patients, of recurrent loops of myelin, called outfoldings (De Sandre-Giovannoli, Delague et al. 2005, Stendel, Roos et al. 2007, Fabrizi, Taioli et al. 2009, Arai, Hayashi et al. 2013, Boubaker, Hsairi-Guidara et al. 2013) These rare myelination abnormalities, have also been reported in other autosomal recessive demyelinating subtypes of Charcot-Marie-Tooth, i.e. CMT4B1 (MIM 601382), CMT4B2 (MIM 604563), and CMT4F (MIM 614895), linked to mutations respectively in *MTMR2* (MIM 603557), *SBF2*/*MTMR13* (MIM 607697), and *PRX* (MIM 605725), suggesting common pathophysiological mechanisms, originating from an initial SC defect (Guilbot and Delague 2001, Bolis, Coviello et al. 2005, Robinson, Niesman et al. 2008). Here, we sought to investigate these mechanisms, by using both *in vitro* and *in vivo* models of CMT4H disease. Indeed, we generated a conditional knock-out mouse model, with specific ablation of *Fgd4/FRABIN* in SCs, that we call *Fgd4*^*SC-/-*^, from which we derived our *in vitro* myelin model based on Dorsal Root Ganglia/Schwann Cells (DRG/SC) cocultures. We showed that the loss of Fgd4/FRABIN in SCs was sufficient to reproduce, both *in vitro* and *in vivo*, the characteristic myelin defects (i.e outfoldings) observed in the nerves of CMT4H patients. *In vivo*, we detected the presence of those myelin abnormalities in the sciatic nerves of *Fgd4*^*SC-/-*^ animals, as early as 3 months of age, and showed that their proportion progressively increase over time. Our results agree with previous data observed in another CMT4H mouse model, using a different promoter (Dhh) to drive Cre recombinase expression in SCs (Horn, Baumann et al. 2012). In our *Fgd4*^*SC-/-*^model, we also provided, for the first time, evidence of late axonal degeneration reflected by late muscle denervation of the gastrocnemius of 12 months old mutant mice, a phenomenon previously described in various de/dysmyelinating CMT (Previtali, Quattrini et al. 2007, Moss, Bopp et al. 2021). Gait test revealed that *Fgd4*^*SC-/-*^ mice display late motor impairment, resulting in the decrease of the pressure exerted by forepaws. These defects may reflect foot misplacement and deformation, as well as distal muscle weakness, and amyotrophy observed in CMT4H patients (De Sandre-Giovannoli, Delague et al. 2005). Overall, our *Fgd4*^*SC-/-*^ mouse model mimics most of the pathogenic features of CMT4H, and is a reliable tool to study the molecular mechanisms underlying CMT4H related aberrant myelination. In this model and the derived *in vitro* myelin model, we showed that the observed myelination defects were connected to a dysregulation of the axo-glial NRG1 type-III/ErbB2/3 pathway and of the downstream effectors P-AKT and mTOR. Notably, we noticed a strong upregulation of P-ErbB2, P-AKT and mTOR in the *Fgd4*^*SC-/-*^ cocultures and in the sciatic nerves of the full knock-out *Fgd4*^*-/-*^ mice. In the PNS, it is known that the amount of axonal NRG1 type III, and its downstream signaling via the activation of the ErbB2/3 receptors, determines the thickness of the myelin sheath (Michailov, Sereda et al. 2004, Taveggia, Zanazzi et al. 2005). Previous work from Bolino et al. (Bolino, Piguet et al. 2016) in models of aberrant focal hypermyelination, suggested that the upregulation of this signaling pathway is likely to contribute to the hypermyelination process, including the generation of aberrant loops of myelin. Also, several studies reported that myelin outfoldings might arise from overactivation of PI3K, AKT and mTOR kinases (Goebbels, Oltrogge et al. 2012, Bolino, Piguet et al. 2016, Domenech-Estevez, Baloui et al. 2016, Figlia, Norrmen et al. 2017). Indeed constitutive activation of AKT or mTORC1 in SCs results in increased thickness and aberrant myelin (Domenech-Estevez, Baloui et al. 2016, Beirowski, Wong et al. 2017, Figlia, Gerber et al. 2018) and the use of the mTORC1 inhibitor, rapamycin, ameliorates myelin outfoldings observed in a DRG/SC coculture model of CMT4B1 (Guerrero-Valero, Grandi et al. 2021).

Moreover, AKT and mTOR levels of expression, as well as activation, are known to be regulated following NRG1type III/ErbB2/3 interaction (Newbern and Birchmeier 2010). In our case, we noticed a significant increase in protein levels of ErbB2 and P-ErbB2 but not of NRG1 type III, suggesting that the loss of Fgd4/FRABIN in SCs may regulate specifically ErbB2 levels rather than NRG1 type III itself. In line with these results, transcriptomic analysis performed on our *Fgd4*^*SC-/-*^ coculture model did not reveal any changes of expression in the NRG1 type III/ErbB2/3 signaling, which suggests a regulation of ErbB2 by FRABIN at the protein level. Interestingly, emerging evidence indicates that ErbB receptor signaling is regulated by endocytic trafficking which controls its degradation or recycling towards the plasma membrane (Sorkin and Goh 2008) (Tomas, Futter et al. 2014, Lee, Chin et al. 2017).

Previous work by Horn and colleagues (Horn, Baumann et al. 2012) have shown that the downregulation of Fgd4/FRABIN in a rat SC line, impairs the endocytosis of the Transferrin receptor (Tfrc). Here, by performing transferrin uptake assays, we showed that the downregulation of Fgd4/FRABIN in primary SC did not affect the basal level of Tfrc ‘s endocytosis but rather promoted its recycling back to the membrane. In *Fgd4*^*SC-/-*^ cocultures, we also observed an upregulation of the small GTPase RAB11, which regulates the trafficking of cargoes to the endocytic recycling compartment for direct recycling to the plasma membrane (Redpath, Betzler et al. 2020). The fate and sorting of cargoes packaged in vesicles of the endo-lysosomal system relies greatly on heterogeneity of vesicles’ membranes, variously enriched in different phosphoinositides that help to shape compartmental identity (Cullen and Carlton 2012). Importantly, FGD4/FRABIN has two PH domains with binding specificity to PI(3,4,5)P3, PI(4,5)P2, and PI(3,4)P2 and one FYVE domain with binding specificity to PI(3)P (Kim, Ikeda et al. 2002, Delague, Jacquier et al. 2007). Our data support the fact that FRABIN, through binding to PIs, regulates the endocytic trafficking of cargo proteins, such as ErbB receptors. Consequently,the enhancement of NRG1 type III/ ErbB2/3 signaling observed in our models could be due to accelerated recycling of the ErbB2 receptor towards the SC plasma membrane rather than to an increased NRG1 type III production. One additional result obtained in our study greatly strengthened this hypothesis. Indeed, by Yeast Two Hybrid experiments, we identified, for the first time, that the Sorting Nexin 3 (SNX3) protein is a new partner of FRABIN in the PNS. Most interestingly, we observed that SNX3 is strongly upregulated in conditions of loss of Fgd4/FRABIN, and conversely, that downregulation of *Snx3* using shRNA, reduces aberrant myelination in *Fgd4*^*SC-/-*^ cocultures. SNX3 belongs to the Sorting nexins (Snxs) family of phosphoinositide-binding proteins and has been shown to be associated with early endosomes through its PX domain. Interestingly, modulation of SNX3 expression has been reported to alter the delivery of Tfrc to recycling endosomes (Xu, Hortsman et al. 2001, Chen, Garcia-Santos et al. 2013).

Overall, our results pointed to defective endosomal recycling as, one main cause of the aberrant myelination associated with the loss of Fgd4/FRABIN, since its blocking using Endosidin-2 decreased the proportion of abnormal myelinated segments in *Fgd4*^*SC-/-*^ cocultures. Interestingly, among others CMT4 related genes, *SH3TC2* (CMT4C) and *MTMR2* (CMT4B1) have been also reported to regulate endocytic trafficking and are connected to defective NRG1 type III/ErbB2/3 signaling (Bolis, Coviello et al. 2009, Lee, Kim et al. 2010, Gouttenoire, Lupo et al. 2013, Bolino, Piguet et al. 2016), which suggests defective endocytic trafficking as a key common pathogenic mechanism for these forms of CMT. Previous work in models of CMT4B2, another neuropathy with focal hypermyelination(Bolino, Piguet et al. 2016), showed that treatment with Niacin efficiently rescued myelination. Nicotinic acid/Niacin, is a compound known to enhance TACE activity (Chen, Cui et al. 2007) and hence negatively regulates NRG1 type-III/ErbB2/3 signaling (Bolino, Piguet et al. 2016). Indeed, BACE-1 and TACE are known to cleave NRG1 type III at distinct cleavage sites, thereby regulating myelination either positively for BACE-1 or negatively for TACE (Hu, Hicks et al. 2006, La Marca, Cerri et al. 2011). Since our observations showed that the dysregulation of the NRG1 type-III/ErbB2/3 signaling pathway was a major contributor to aberrant myelination in CMT4H, we used Niacin to assess its capacity to rescue myelin outfoldings in our models. Remarkably, the use of Niacin improved myelination of both *Fgd4*^*SC-/-*^ cocultures and animals’sciatic nerves. We chose to start Niacin treatment at 1 month of age, i.e., before the appearance of myelin defects in the nerves of *Fgd4*^*SC-/-*^ mice. Horn and colleagues (Horn, Baumann et al. 2012) showed that FRABIN seems involved in myelin maintenance rather than in myelination initiation since the induction of its deletion from 2-month-old was sufficient to produce myelin abnormalities comparable to the *Fgd4*^*SC-/-*^ animals. Based on this assumption, we could therefore anticipate that counteracting FRABIN’s loss-associated defects (i.e., NRG1 type III/ErbB2/3 pathway) using Niacin at a later time-point could provide benefits for CMT4H pathology. Overall, we have demonstrated that the specific loss of FGD4/FRABIN in SCs was sufficient to induce abnormal myelination (i.e outfoldings) like the ones observed in the nerves of patients affected with CMT4H and that this aberrant myelination was accompanied by an upregulation of the NRG1 type III/ ErbB2/3 pathway, as well as of downstream effectors (AKT/mTOR). Our results showed that targeting NRG1 type-III/ErbB2/3 signaling using drugs based on nicotinic acid/niacin, could be an effective therapeutic perspective for demyelinating CMT subtypes associated with focal hypermyelination. In addition, we provided new insights into the pathophysiological mechanisms underlying CMT4H, by demonstrating that FRABIN regulated endocytic trafficking, in particular of the ErbB2 receptor. Our results open new avenues for a better understanding and treatment of CMT4H as well as for other hypermyelinating neuropathies.

## Acknowledgements

The electron microscopy experiments were performed on the **PiCSL-FBI core facilty (IBDM, AMU-Marseille)**, member of the France-BioImaging national research infrastructure. Sequencing and bioinformatics analysis were performed by the **Genomics and Bioinformatics facility (GBiM)** from the U 1251/Marseille Medical Genetics lab. This research work was supported by the French Association against Myopathies (Association Française contre les Myopathies, Grants # MNH-Decrypt, # TRIM-RD and #MoTharD).

The Authors declare that there is no conflict of interest.

## Authors contributions

NBM, LEB, PR and VD designed the experiments; NBM, LEB, NT, PQ, CE, AG performed the experiments; NBM, LEB, PR, NL and VD analyzed the data; NBM, LEB and VD wrote the paper. PR, YP and MB edited and reviewed the manuscript.

