## Supplemental Information for "Imbalance of Neuregulin1-ErbB2/3 signaling underlies altered myelin homeostasis in models of Charcot-Marie-Tooth disease type 4H"

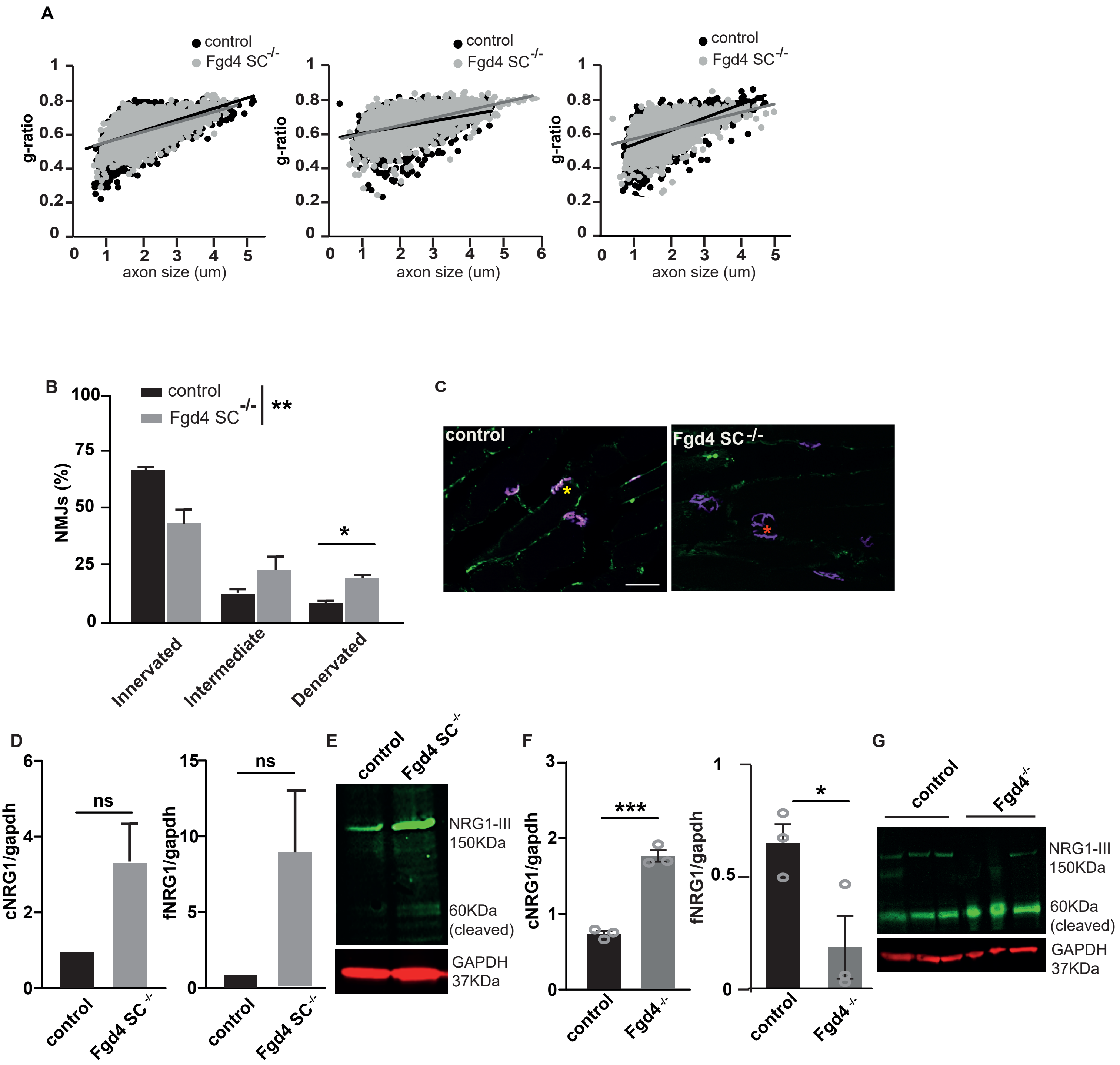
A
control
Fgd4 SC-/-
g-ratio
axon size (um)
B
control
Fgd4 SC-/-
**
NMJs (%)
*
Innervated
Intermediate
Denervated
C
control
Fgd4 SC-/-
D
cNRG1/gapdh
fNRG1/gapdh
ns
control
Fgd4 SC-/-
E
control
Fgd4 SC-/-
NRG1-III 150KDa
60KDa (cleaved)
GAPDH 37KDa
F
cNRG1/gapdh
***
fNRG1/gapdh
*
control
Fgd4-/-
G
control
Fgd4-/-
NRG1-III 150KDa
60KDa (cleaved)
GAPDH 37KDa

**Figure Sup 1**

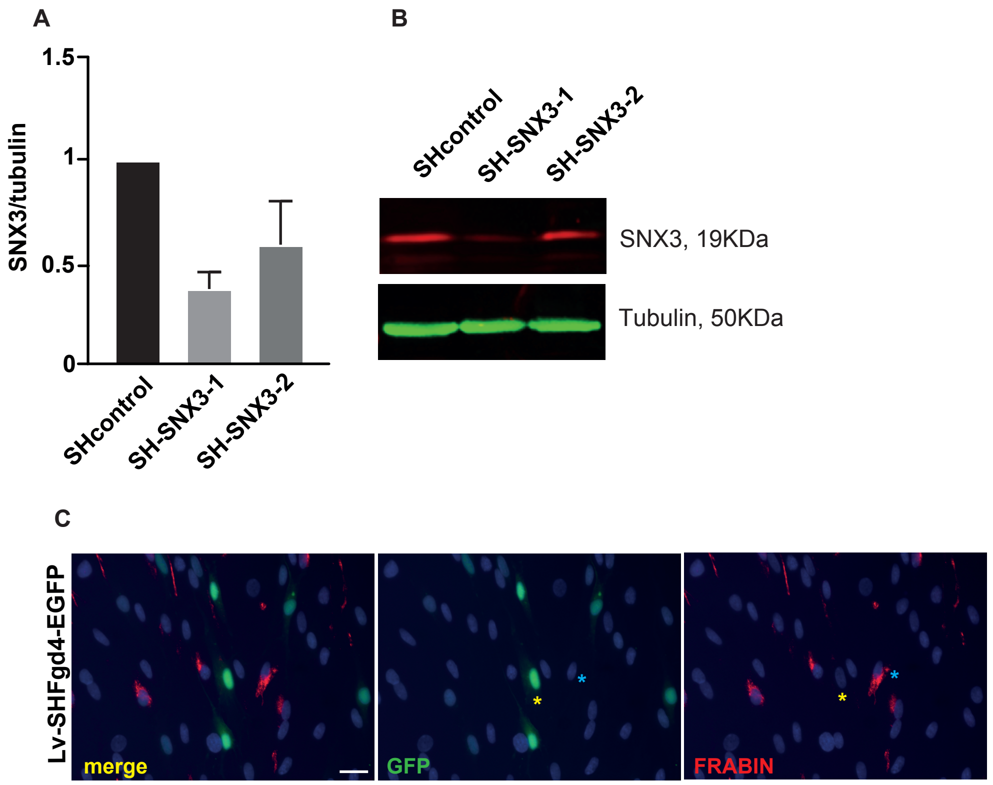
A
1.5
1
0.5
0
SNX3/tubulin
SHcontrol
SH-SNX3-1
SH-SNX3-2
B
SHcontrol
SH-SNX3-1
SH-SNX3-2
SNX3, 19KDa
Tubulin, 50KDa
C
Lv-SHFgd4-EGFP
merge
GFP
FRABIN

**Figure sup 2**

**Figure Sup1.** ***Fgd4***$^{SC-/-}$ **animals display no defects of myelination thickness, but a late muscle denervation** **(A)** g-ratio analysis revealed no statistical difference in myelin thickness in the sciatic nerves of 3, 6 and 18 mo WT and *Fgd4*$^{SC-/-}$ mice. A total of 500-1000 axons of diameter between 0.5 and 6 µm were analysed (n=3 animals per genotype). Data are represented as scatter plots of individual axons as a function of their respective diameters determined at 3, 6 and 18 months old. Each point corresponds to one fiber (gray points: *Fgd4*$^{SC-/-}$ animals; black points: WT animals). (**B-C**) Late muscle denervation observed in the gastrocnemius of *Fgd4*$^{SC-/-}$ animals. Level of innervation of gastrocnemius muscle evaluated by the colocalization of the neurofilament marker (NF-M) and the acetylcholine receptor marker $\alpha$-bungarotoxin in 12 months old WT (n=3) and *Fgd4*$^{SC-/-}$ (n=3) animals. Yellow asterisk indicate innervated NMJ, red asterisk indicate denervated NMJ. Scale bar: 50 µm. Statistical analysis: two-way repeated-measures ANOVA (genotype*type of NMJs) with Sidak post-hoc test. Two-way ANOVA revealed a significant difference on the proportion of the type of NMJs between WT and *Fgd4*$^{SC-/-}$ conditions (p=0.007). Sidak multiple comparison test show a significant increase in the proportion of denervated NMJs in *Fgd4*$^{SC-/-}$ compared to WT (*p*:0,01). (**D-E**) Levels of expression of cleaved and full-length Neuregulin 1-type III (named respectively cNRG1 and fNRG1) were assessed by western-blot analysis in *Fgd4*$^{SC-/-}$ cocultures compared to control. (**D**) Data are expressed as mean ± sem (n=3-4 cocultures). Statistical analysis: unpaired Student's t-test. (**E**) Western blot pictures illustrating the expression of the markers described in (D). (**F-G**) Levels of expression of cNRG1 and fNRG1 were assessed by western-blot analysis in the sciatic nerves of *Fgd4*$^{-/-}$ mice compared to WT mice. (**F**) Data are expressed as mean ± sem (n=3 animals per genotype). Statistical analysis: unpaired Student's t-test. (**G**) Western blot pictures illustrating the expression of the markers described in (E). * p<0.05, ** p<0.01, ***p<0.001.

**Figure Sup2. Effective knock-down of Sn3 and Fgd4/FRABIN in vitro**. (**A-B**) Lentivirus (Lv) expressing shRNA targeting Snx3 lead to an efficient knock-down of Snx3 in primary rat SCs. Primary SCs were infected 1 day after plating with either Lv-SHcontrol, LV-SH-SNX3-1 or 2, and harvested 7 days post-infection. Level of expression of SNX3 in those conditions was evaluated by western-blot. (**C**) Knock-down of *Fgd4/FRABIN* in primary SCs following Lv-shRNA targeting *Fgd4* and expressing GFP tag SHFgd4. Primary SCs were infected 1 day after plating and fixed 3 days post-infection. Infected cells were identified by GFP expression. Expression levels of FRABIN were evaluated by immunofluorescence (in red). In contrast to non-infected cells (GFP negative, blue asterisks) which express FRABIN, infected cells (GFP positive cells, yellow asterisks) are negative for FRABIN.
